# Calibrated ADAMTS13 inhibition prevents cardiovascular shear coagulopathy

**DOI:** 10.64898/2026.08.03.742456

**Authors:** Kenki Saito, Noriyoshi Isozumi, Yasuyuki Shiraishi, Yoshikazu Hattori, Kazuya Sakai, Tomoya Ueda, Atsushi Hamamura, Ryota Imamura, Michinori Kayashima, Kotaro Nakano, Tomoyuki Yambe, Hiroyuki Kumeta, Ryouta Shigehisa, Masashi Mori, Tomohiro Imamura, Mari Nakanishi, Masayuki Oda, Shingo Kanemura, Masaki Okumura, Tatsuya Niwa, Anne Martel, Lionel Porcar, Ken Morishima, Aya Okuda, Masaaki Sugiyama, Miki Takatsuka, Nozomi Tomimatsu, Tomohide Saio, Shungo Hikoso, Eiichiro Mori, Masanori Matsumoto

## Abstract

Mechanical circulatory support essential for managing severe heart failure frequently triggers bleeding complications, driven by shear stress-induced over-proteolysis of von Willebrand factor (VWF) by ADAMTS13. Inhibiting ADAMTS13 presents a rationale to treat this condition, known as acquired von Willebrand syndrome (AVWS). However, conventional therapeutic strategies remain limited due to the risk of triggering thrombotic thrombocytopenic purpura. Here we show that HA10, a humanized anti-ADAMTS13 antibody, preserves a residual level of ADAMTS13 activity above the thrombosis-associated threshold. Multimodal structural and biophysical analyses—including NMR, SAXS, and SANS—revealed that HA10 bound to the disintegrin-like domain of ADAMTS13, dynamically competing with VWF while leaving 10– 20% residual enzymatic activity. The therapeutic efficacy and safety of HA10 were verified in non-human primate models of AVWS. Our findings establish a novel paradigm of enzymatic calibration rather than complete blockade, offering a mechanistically targeted and safe therapeutic approach for cardiovascular bleeding.

## Main

Mechanical circulatory support (MCS) technologies, including left ventricular assist devices (LVAD), extracorporeal membrane oxygenation (ECMO), and catheter-based micro-axial flow pump (Impella) support, are essential modalities for managing severe heart failure. Despite their life-saving benefits, these devices expose blood to pathological fluid shear stress, frequently triggering bleeding complications^1–4^. A primary driver of this mechanical coagulopathy is acquired von Willebrand syndrome (AVWS)^5–7^, wherein shear-induced conformational extension of circulating von Willebrand factor (VWF) accelerates its over-proteolysis by the metalloproteinase ADAMTS13 (Fig. 1a)^8–10^.

**Fig. 1:**
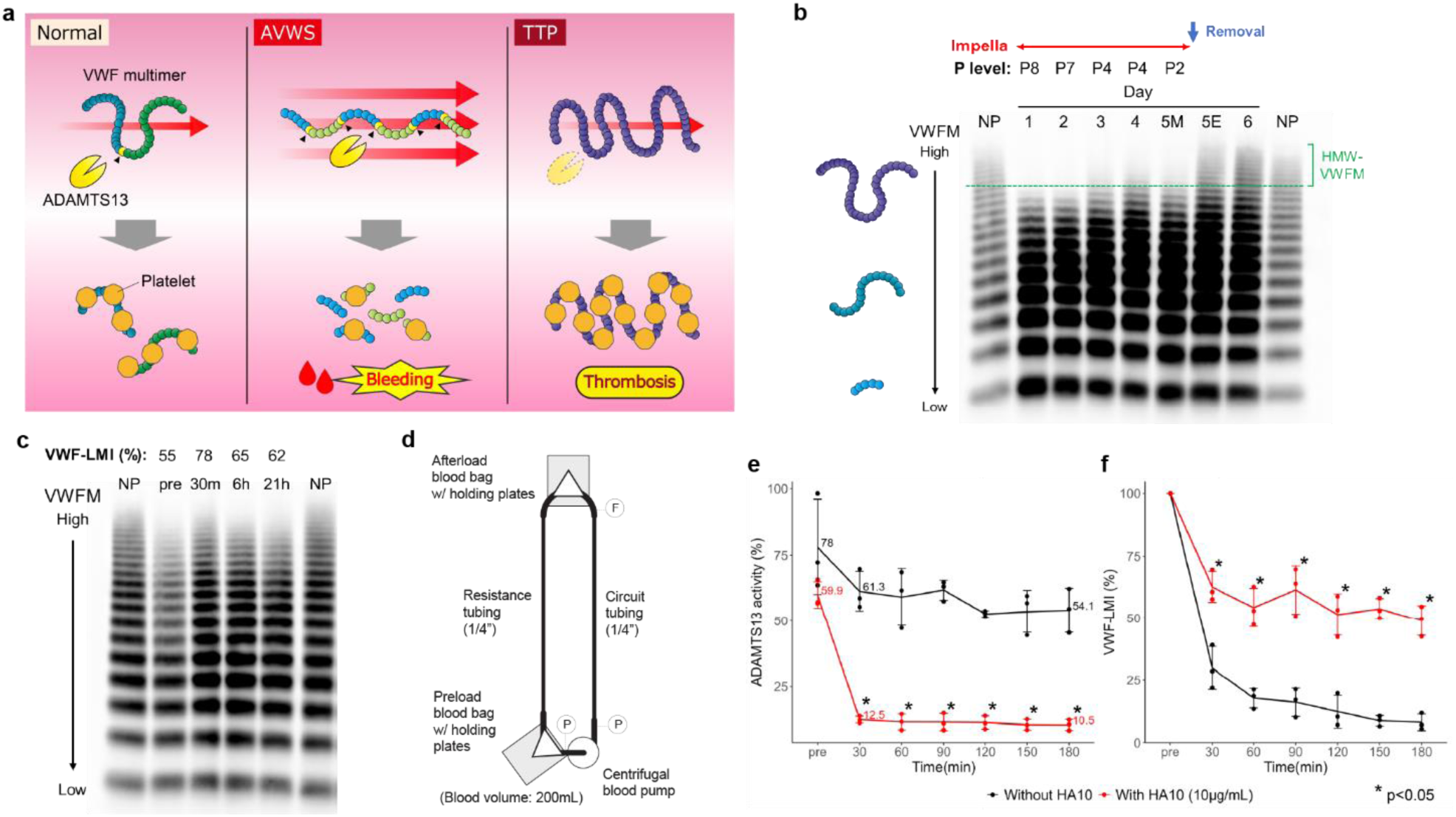
HA10 blocks degradation of VWF. **a**, Schematic of normal hemostasis, AVWS, and TTP, illustrating changes in VWF multimers, ADAMTS13 activity, and platelet thrombus formation. **b**, Temporal changes in VWF multimers in an illustrative patient, in whom Impella was administered as therapy for acute heart failure associated subacute myocardial infarction. Impella support was continued for 5 days and discontinued between Day 5M (morning) and Day 5E (evening). HMW-VWFM disappeared during the treatment. HMW-VWFM gradually restored following the decline in the Impella support levels (P levels, index range: P1-P9) and exhibited substantial recovery immediately after the removal of Impella. **c**, Temporal changes in VWF multimers during the recombinant VWF (vonicog alfa) infusion test conducted as a preoperative evaluation for AVWS associated with severe mitral regurgitation. Even after the administration of 1300 IU (approximately 30 IU/kg), HMW-VWFM did not fully restore, with VWF-LMI remaining below 80%, a condition defined as loss of large VWF multimers. **d**, Schematic illustration of an ex vivo human AVWS model using a PCPS circuit. **e**–**f**, Temporal changes in ADAMTS13 activity (**e**) and VWF-LMI (**f**) in the *in vitro* PCPS perfusion experiments using 200 mL of healthy donor blood. Error bars represent mean ± SD. *p < 0.05. Abbreviations: VWF, von Willebrand factor; HMW-VWFM, High-molecular-weight VWF multimers; PCPS, percutaneous cardio-pulmonary support; VWF-LMI, VWF large multimer index.

This aberrant shear-responsive proteolysis is a hallmark not only of device-implanted patients but also of diverse cardiovascular pathologies, such as aortic stenosis (Heyde syndrome)^11^ and hypertrophic obstructive cardiomyopathy^12^. Addressing this molecular mechanism remains a major therapeutic challenge; while shutting down ADAMTS13 proteolysis is a rational approach to prevent VWF degradation^13^, a complete enzymatic blockade poses a risk of triggering microvascular thrombosis, as seen in thrombotic thrombocytopenic purpura (TTP) (Fig. 1a)^14,15^.

Here we show that HA10, a humanized anti-ADAMTS13 antibody^16^, resolves this trade-off by acting as a molecular rheostat that calibrates VWF proteolysis within a safe physiological window, preventing AVWS without provoking thrombotic complications in non-human primate models. By integrating multimodal biophysical and structural approaches—including nuclear magnetic resonance (NMR), analytical ultracentrifugation (AUC), small-angle X-ray scattering (SAXS), and small-angle neutron scattering (SANS)—we elucidate the dynamic structural basis of this partial inhibition, demonstrating how HA10 couples with the disintegrin-like domain and responds to conformational transitions of ADAMTS13.

## Results

### An unmet clinical need for calibrating shear-induced VWF proteolysis

VWF is a large multimeric glycoprotein that functions as a vascular mechanosensor, undergoing profound conformational transitions in response to hydrodynamic shear forces^17,18^. Under physiological flow, VWF adopts a globular state that sequesters the crucial proteolytic cleavage site within its A2 domain^19^. However, pathological shear stress—such as that generated by MCS devices forces VWF into an extended conformation, exposing the Tyr1605–Met1606 peptide bond to accelerated cleavages by ADAMTS13^20–23^. This aberrant structural transition leads to a depletion of high-molecular-weight VWF multimers (HMW-VWFM), which are indispensable for high-shear platelet adhesion, clinically manifesting as AVWS and bleeding complications (Fig. 1a)^18,24,25^.

To define this mechanical coagulopathy in a clinical setting, we monitored a patient undergoing Impella support. The circulating HMW-VWFM was obliterated during device operation but recovered upon Impella removal, demonstrating the link between mechanical shear and the loss of hemostatic multimers (Fig. 1b). The therapeutic failure of standard approaches was further highlighted in an AVWS case secondary to severe mitral regurgitation, in which an attenuated *in vivo* effect of recombinant VWF concentrate due to its immediate, shear-driven degradation was demonstrated. (Fig. 1c). This rapid destruction is consistent with the results from a recent randomized controlled trial^26^ where prophylactic VWF supplementation failed to prevent hemorrhage in LVAD patients due to a transient prohemostatic window. Collectively, these findings underscore an unmet need, suggesting that rather than replenishing the rapidly degraded substrate, therapeutic strategies should target the catalytic machinery itself to prevent the mechanical destruction of HMW-VWFM.

To test whether modulating ADAMTS13 proteolysis could rescue HMW-VWFM under pathological flow, we evaluated HA10—a recently generated humanized anti-ADAMTS13 antibody— in an *ex vivo* human AVWS model using a percutaneous cardio-pulmonary support (PCPS) circuit (Fig. 1d). Targeting ADAMTS13 with GLP-grade HA10 successfully suppressed excessive enzymatic activity (Fig. 1e), thereby rescuing the VWF large multimer index (VWF-LMI)^27^ from shear-induced reduction (Fig. 1f). These results demonstrate that HA10-mediated calibration of ADAMTS13 activity represents a potent and rational therapeutic strategy to counteract AVWS.

### Structural basis for HA10-mediated partial inhibition *via* ADAMTS13 disintegrin-like domain

To elucidate the structural basis by which HA10 calibrates VWF cleavage, we mapped its binding interface on ADAMTS13 using an integrated multimodal biophysical approach. The original mouse monoclonal antibody A10 (mA10) was previously shown to recognize the disintegrin-like (D) domain of human ADAMTS13^28^, yet the precise atomic determinants remained elusive. We successfully prepared properly folded and disulfide-bonded recombinant ADAMTS13 D-domain (ADAMTS13D) (Extended Data Fig. 1) and subjected it to solution NMR spectroscopy for sequence-specific backbone resonance assignments (Extended Data Fig. 2).

NMR analysis following the addition of mA10 and HA10 identified a contiguous epitope cluster. This binding interface comprises residues Q310, C332, H338, T339, R349, and C371 (Fig. 2a,b). The functional relevance of this interface was further validated through site-directed mutagenesis and biochemical binding assays, confirming the critical involvement of Q310, H338, T339, and R349 (Fig. 2c). To visualize the solution structure of the complex, we performed small-angle X-ray and neutron scattering (SAXS/SANS) combined with analytical ultracentrifugation (AUC) (Extended Data Figs. 3 and 4). Computational docking using these data generated a high-resolution structural model of the ADAMTS13–HA10 complex (Fig. 2d). Notably, the model demonstrates that HA10 directly covers the highly conserved exosite required for VWF substrate recognition within the D domain^29,30^.

**Fig. 2:**
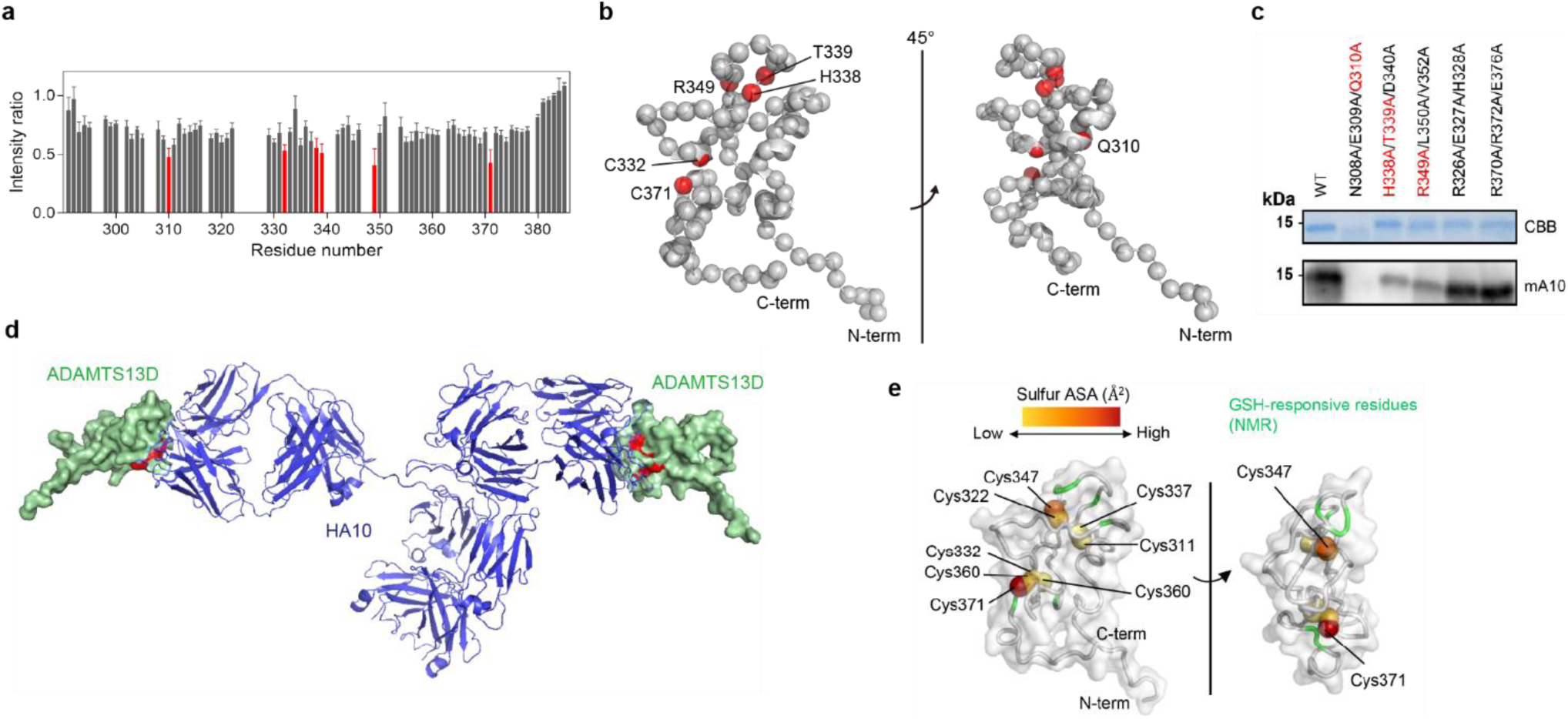
HA10 recognizes disintegrin-like domain of ADAMTS13. **a**, Residue-specific ^1^H-^15^N NMR signal intensity changes of ^15^N-labeled ADAMTS13D upon HA10 binding. The intensity ratio represents the signal intensity in the HA10-bound state relative to the free state. Residues showing marked intensity changes (below the mean−SD) are indicated by red bars. **b**, Ribbon model of ADAMTS13D. Residues showing marked intensity changes in panel **a** are highlighted here in red and labeled by amino acid type and residue number. **c**, Mutational validation of the HA10-binding interface identified by NMR. Wild-type (WT) and mutant ADAMTS13D were analyzed using mA10-binding assays. NMR-identified candidate residues showing reduced mA10 binding upon mutation are highlighted in red. **d**, HA10-ADAMTS13D complex model based on the SAXS data (Extended Data Fig. 4). Blue and green parts show HA10 and ADAMTS13D, respectively. The VWF-binding exosite within ADAMTS13D is highlighted in red. Details of structural modeling are described in the Methods section. **e**, Structural mapping of cysteine accessibility and GSH-responsive residues of ADAMTS13D. Sulfur atoms of cysteine residues involved in disulfide bonds are represented as spheres colored according to their accessible surface area (ASA), with higher ASA values shown in red and lower ASA values in yellow. The C322–C347 and C360–C371 disulfide bonds exhibited greater solvent accessibility (ASA values of 0.86 and 3.24 Å^2^, and 0.34 and 8.51 Å^2^, respectively) than the buried C311–C337 and C332–C366 disulfide bonds (ASA = 0 Å^2^). Residues showing marked intensity changes upon GSH addition, identified by solution NMR spectroscopy, are highlighted in green.

### Conformational plasticity of the C371 residue dictates incomplete complex formation

A central mystery of HA10 is why a 10–20% residual enzymatic activity persists even at saturating concentrations (Fig. 1e and Extended Data Fig. 5). We hypothesized that the internal disulfide bond (S-S) network within ADAMTS13D governs this structural calibration. Systematic serine-substitutions of the eight cysteines (forming four S-S bonds) revealed a striking hierarchy: disrupting C311–C337 (SS1) or C332–C366 (SS3) destabilized the domain, whereas mutating C322– C347 (SS2) or C360–C371 (SS4) exerted remarkably mild effects (Extended Data Fig. 6a,b). This indicates that while SS1 and SS3 form the rigid, structurally buried core, SS2 and SS4 possess high local flexibility.

Remarkably, solvent-accessible surface area (ASA) calculations highlighted that the sulfur atom of C371 (within SS4) is exceptionally exposed to the bulk solvent (Fig. 2f and Extended Data Fig. 6c). This solvent accessibility was corroborated by NMR analysis in the presence of glutathione (GSH) (Fig. 2f). These findings suggest that the conformational plasticity around C371 contributes to the incomplete inhibition of ADAMTS13 by HA10, thereby providing a structural basis for the safe, calibrated inhibition of VWF proteolysis.

### Competitive displacement of VWF from the ADAMTS13 exosite underlies partial inhibition

While the catalytic metalloprotease (M) domain directly cleaves the VWF A2 domain^31^, HA10 achieves its inhibitory effect by targeting the non-catalytic disintegrin-like (D) domain^28^. This raised a fundamental question: why does an antibody targeting the D domain, rather than the catalytic M domain, effectively modulate proteolytic activity? To address this, we investigated how HA10 interferes with the physical assembly of the VWF–ADAMTS13 complex.

Among the epitope residues identified for HA10, the R349, L350, and V352 triplet constitutes a highly conserved exosite known to anchor the unfolded VWF A2 domain (Extended Data Fig. 7)^29,30^. To directly visualize this molecular interference, we performed solution NMR spectroscopy using isotopically labeled VWF73 (residues D1596–R1668 of VWF) (Fig. 3a and Extended Data Fig. 8)^32^. Upon incubating VWF73 with ADAMTS13D, we observed pronounced chemical shift perturbations within the Y1605–G1629 region of VWF73, confirming its direct engagement with the D-domain exosite (Fig. 3b). Remarkably, the subsequent addition of HA10 completely displaced VWF73 from ADAMTS13D, restoring the free-state NMR spectra of VWF73 (Fig. 3c).

**Fig. 3:**
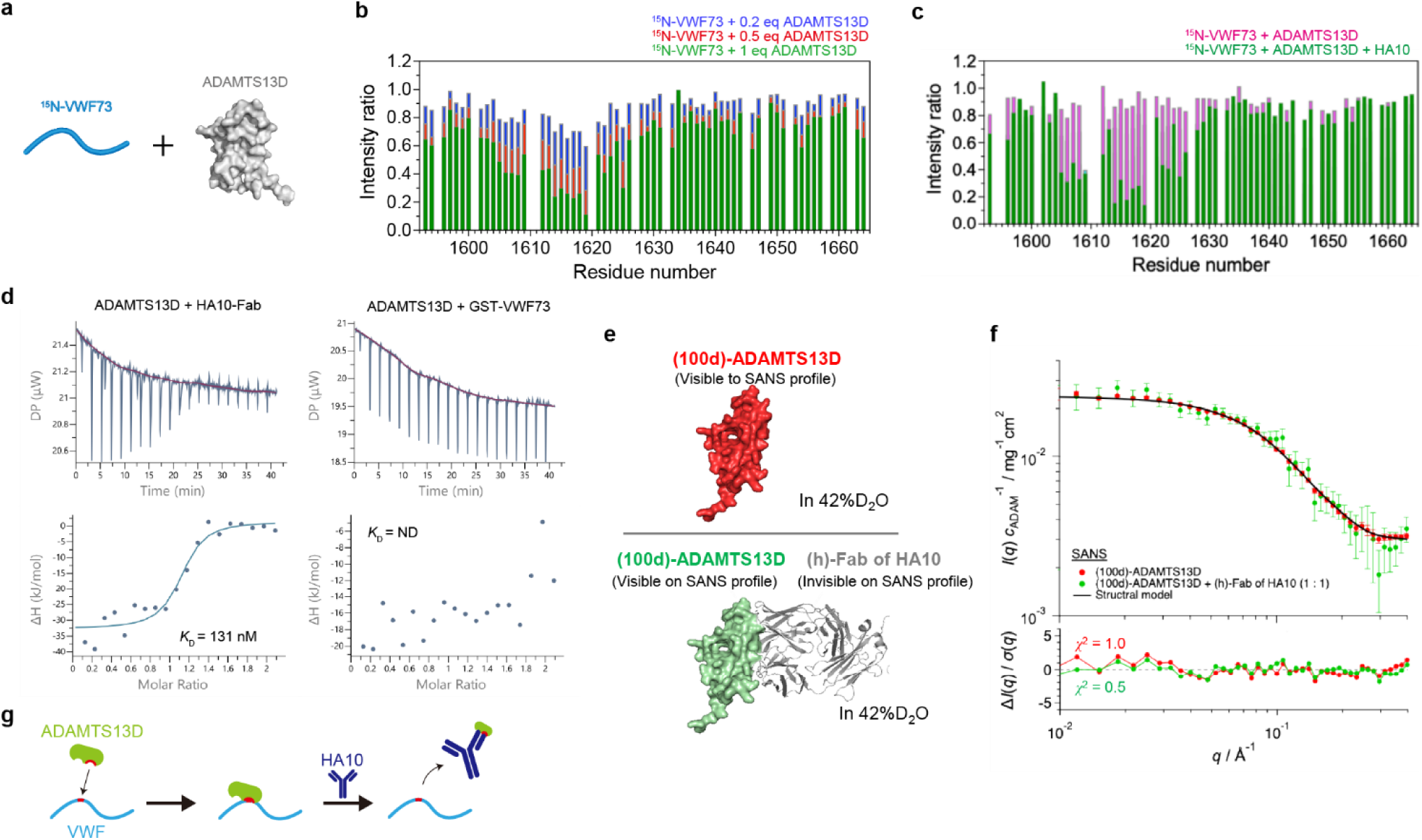
HA10 competes with VWF binding to ADAMTS13. **a**, Schematic of the ^1^H-^15^N NMR experiment using ^15^N-labeled VWF73 (shown in cyan) and ADAMTS13D (shown in gray). **b**, Residue-specific ^1^H-^15^N NMR signal intensity changes of ^15^N-labeled VWF73 upon titration with ADAMTS13D at 0.2, 0.5, and 1.0 eq (blue, red, and green bars, respectively). The intensity ratio represents the signal intensity relative to free ^15^N-labeled VWF73. **c**, Residue-specific signal intensity changes of ^15^N-labeled VWF73 in the presence of ADAMTS13D (green bars) and following the addition of HA10 (magenta bars). **d**, ITC measurements of ADAMTS13D binding to HA10-Fab and GST-VWF73. The dissociation constant (*K*_D_) was determined for HA10-Fab but not for GST-VWF73. ND: not determined because only dilution heat was observed. **e**, Structural model of 100%-deuterated (100d)-ADAMTS13D (upper) and (100d)- ADAMTS13D + hydrogenated (h)-Fab complex (lower). **f**, Red and green circles represent the SANS profile of (100d)-ADAMTS13D in 42%D_2_O and (100d)-ADAMTS13D + (h)-Fab of HA10 complex in 42%D_2_O, respectively. The scattering intensities are normalized by concentration of (100d)- ADAMTS13D (*c*_ADAM_). Because the scattering length density of (h)-Fab matches that of 42% D_2_O, (h)-Fab is invisible on SANS profile. The agreement between the two scattering profiles means that the partial structure of ADAMTS13D in the complex is identical to ADAMTS13D alone in solution. Black solid line represents the scattering profile calculated for ADAMTS13D. The residuals between the experimental and calculated scattering profiles are shown in the lower panels. **g**, Schematic illustrating the proposed mechanism of HA10-mediated competitive inhibition of ADAMTS13-VWF binding without inducing TTP.

To quantify the energetic basis of this molecular displacement, we conducted isothermal titration calorimetry (ITC) analysis (Fig. 3d). Consistent with the NMR competition experiments, HA10 bound ADAMTS13D with high affinity (*K*_D_ = 131 nM). In contrast, VWF73 binding was not detectable by ITC under comparable experimental conditions, with only dilution heat observed, suggesting a markedly weaker interaction than HA10 under these experimental conditions. These thermodynamic data indicate that HA10 does not sterically lock the active site; instead, it establishes a delicate dynamic equilibrium with the substrate. Supporting this model, SANS analysis showed that HA10 binding does not induce detectable conformational changes in ADAMTS13D (Fig. 3e,f). Under pathological shear stress, the high mechanical force applied to multimeric VWF allows it to transiently outcompete the antibody at the exosite, providing a mechanistic explanation for the 10–20% residual enzymatic activity. Collectively, these results elucidate how targeting a flexible exosite, rather than the absolute blockade of the catalytic center, enables the calibrated regulation of VWF proteolysis (Fig. 3g).

### Primate-specific exosite recognition enables targeted intervention in a non-human primate AVWS model

To evaluate the translational potential of HA10, we first characterized its species cross-reactivity using an *in vitro* high-shear syringe model^33^. HA10 successfully prevented shear-induced HMW-VWFM degradation in human and cynomolgus monkey (*Macaca fascicularis*) plasma, but exhibited no inhibitory efficacy in bovine, porcine, or caprine systems (Fig. 4a). Interspecies sequence alignment, coupled with site-directed mutagenesis, revealed that residues S306 and N308 within the D domain dictate this strict primate-specificity (Fig. 4b,c, and Extended Data Fig. 7). This evolutionary divergence mandated the use of a non-human primate model to predict clinical efficacy and safety. Crucially, in a cynomolgus monkey AVWS model utilizing an extracorporeal PCPS circuit (Extended Data Fig. 9a), administration of HA10 robustly rescued the shear-induced depletion of circulating HMW-VWFM (Extended Data Fig. 9b), establishing its potent *in vivo* therapeutic efficacy within a primate vascular environment.

**Fig. 4:**
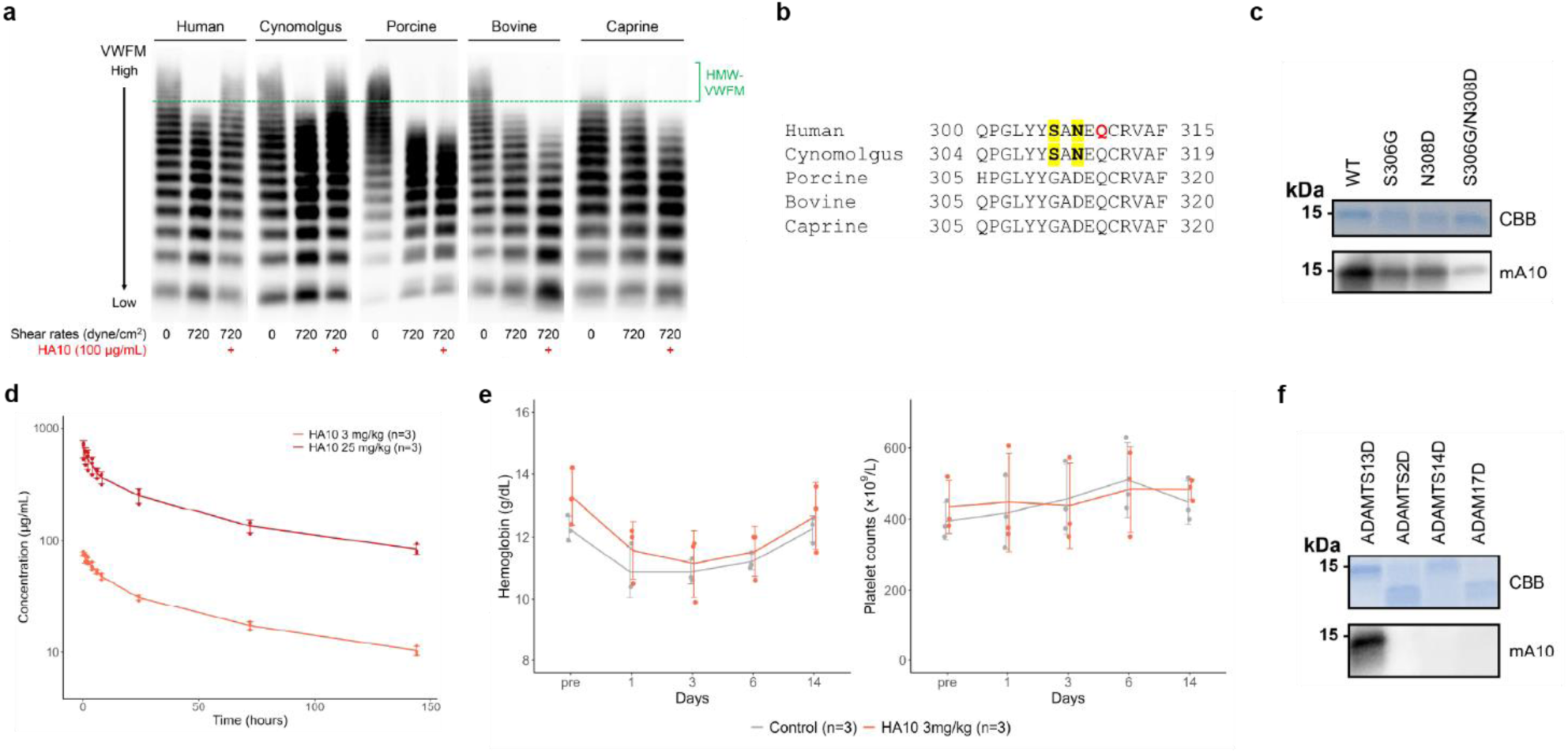
HA10 did not induce TTP-like thrombotic events in cynomolgus monkeys. **a**, VWF multimer analysis of shear-stressed plasma samples in an *in vitro* syringe model as described in the Methods section. **b**, Partial amino acid sequence alignment of ADAMTS13D from different species. Residues conserved in human and cynomolgus, but not in porcine, bovine, or caprine sequences, are highlighted in yellow. The HA10-binding residue Q310, identified in Fig. 2, is shown in red. **c**, mA10-binding assays of ADAMTS13D mutants. Residues S306 and/or N308 in wild-type (WT) human ADAMTS13D were substituted with corresponding porcine, bovine, or caprine residues. **d**, Toxicokinetic profile of HA10 following a single GLP-grade dose in male cynomolgus monkeys. Mean elimination half-lives were 138 h at 3 mg/kg group and 159 h at 25 mg/kg group. Error bars represent mean ± SD. **e**, Temporal changes in hemoglobin and platelet counts over time following the same HA10 administration in the 3 mg/kg (n = 3) and control (n = 3) groups. Error bars represent mean ± SD. *p < 0.05. **f**, mA10-binding assays using the disintegrin-like (D) domains from ADAMTS and ADAM family members. Abbreviations: VWF-LMI, VWF large multimer index.

### *In vivo* safety profiling demonstrates avoidance of TTP-like microvascular thrombosis

A formidable barrier to therapeutic ADAMTS13 inhibition is the risk of microvascular thrombosis, as classically demonstrated by metalloprotease-domain-targeted antibodies that induce fatal TTP phenotypes in baboon models (Fig. 1a)^15^. According to the International Society on Thrombosis and Haemostasis (ISTH) guidelines, a clinical diagnosis of TTP requires depletion of ADAMTS13 activity below the 10% threshold^34,35^. Our *in vitro* profiling confirmed that even at saturating concentrations up to 10 μg/mL, HA10-mediated inhibition strictly plateaued, maintaining 10–20% residual enzymatic activity (Fig. 1e and Extended Data Fig. 5).

To verify that this structural calibration translates to safety *in vivo*, we performed a safety and pharmacokinetic evaluation in cynomolgus monkeys at doses of 3 and 25 mg/kg, with a vehicle control group (n = 3 per group). Single-dose intravenous administration of HA10 at a therapeutic dose (3 mg/kg) or a supratherapeutic dose (25 mg/kg) revealed a pharmacological half-life of 138 and 159 hours (∼6–7 days), respectively (Fig. 4d). Of note, monkeys treated with the therapeutic dose exhibited no clinical, hematological, or histopathological signs of microvascular thrombosis (Fig. 4e). Furthermore, even at the 8-fold supratherapeutic dose (25 mg/kg), the animals showed no overt clinical or histopathological evidence of TTP-like pathology (Extended Data Fig. 9c–g).

These *in vivo* data demonstrate that HA10 mitigates the thrombotic liability associated with complete ADAMTS13 blockade. To evaluate target specificity, cross-reactivity was assessed against representative ADAMTS and ADAM family members. mA10-binding assays revealed no detectable cross-reactivity with the corresponding D domains of these family members, further supporting selective recognition of ADAMTS13D by HA10 (Fig. 4f). By biological design, the dynamic structural constraints of the C371-modulated exosite guarantee that residual ADAMTS13 activity does not drop below the 10% ISTH threshold. Consequently, HA10 achieves an unprecedented therapeutic window, reconciling anti-bleeding efficacy with thrombotic safety in primates.

## Discussion

The clinical management of shear-induced acquired von Willebrand syndrome (AVWS) has long faced a major therapeutic challenge: whether to continuously replenish the depleted substrate or to target the underlying catalytic machinery. A recent randomized controlled PHAM trial^26^, which evaluated a purified VWF concentrate during mechanical circulatory support (MCS), confirmed a rapid kinetic of degradation of VWF concentrate in LVAD setting. This pivotal clinical outcome implies that substrate replenishment is inadequate under pathological fluid dynamics. Instead, therapeutic intervention needs to pivot toward modulating the endogenous shearing enzyme, ADAMTS13.

However, therapeutic inhibition of ADAMTS13 introduces a safety concern. Previous pioneering work demonstrated the feasibility of treating AVWS *via* ADAMTS13 blockade in a calf model^13^, yet the same group also successfully established a primate TTP model by completely ablating ADAMTS13 activity *via* metalloprotease-domain-targeted antibodies in baboons^15^. These contrasting studies highlighted an important issue: complete enzymatic blockade leaves minimum safety margin against microvascular thrombosis. Our study resolves this paradigm by demonstrating that targeting the non-catalytic disintegrin-like (D) domain, rather than the catalytic metalloprotease center, shifts the inhibitory mechanism from steric occlusion to dynamic substrate competition.

This mechanistic divergence provides the structural basis for the unique safety profile of HA10. Our multimodal structural analyses reveal that HA10 competes with the unfolded VWF A2 domain at the D-domain exosite, while the solvent-exposed C371 residue undergoes an open-close conformational transition that limits the affinity of the antibody. Consequently, even at a maximum serum concentration approaching 50 μg/mL on Day 1 following a 3 mg/kg dose, HA10-mediated inhibition plateaus, ensuring that residual ADAMTS13 activity remains above the 10% threshold defined by the latest ISTH diagnostic guidelines^34^. This incomplete, calibrated inhibition allows high-shear-stretched, massive multimeric VWF molecules to transiently outcompete the antibody and undergo essential baseline proteolysis, effectively bypassing the dangerous threshold of TTP development.

The pharmacokinetics of HA10 further underscore its clinical utility. With a primate half-life of approximately 6–7 days, a single 3 mg/kg intravenous dose maintains protective serum levels above the IC50 (∼1 μg/mL) for over a week, without inducing any hematological or histopathological signs of thrombosis. This pharmacological profile enables tailored clinical regimens: a single-shot administration for acute-phase support during temporary Impella or ECMO/PCPS tracking, or a convenient biweekly maintenance regimen for chronic-phase protection in patients with durable LVADs or underlying valvular pathologies such as Heyde syndrome.

In conclusion, our integration of multi-scale structural biology, biophysical competition, and rigorous non-human primate GLP safety profiling establishes a novel therapeutic framework. By substituting absolute enzymatic blockade with molecular calibration, HA10 paves the way for upcoming first-in-human clinical trials, offering a mechanistically targeted, safe, and mechanistically grounded therapeutic strategy for cardiovascular shear-induced coagulopathies.

## Methods

### VWF multimer analysis

VWF multimer analysis was performed as described previously^37^. Large VWF multimers were defined as the 11th peak or higher than the lowest molecular weight fractions in the densitometric analysis using ImageJ software^38^. VWF multimer index was calculated as the ratio of the area under the curve of large multimers in the sample relative to that of the control^27^. Pooled normal plasma was used as the control for patient sample analysis, whereas baseline (pre-experiment) samples were used as the control for the *in vivo* analysis.

### Case presentation

#### A-73-year-old male with Impella treatment

Serial patient plasma samples were collected during and after the Impella treatment, and VWF multimer analysis was performed. This series of analyses was approved by the Nara Medical University ethics committee (approval no. G107).

#### A-81-year-old female with severe mitral regurgitation

A recombinant VWF (vonicog alfa, Takeda, Tokyo, Japan) infusion test was conducted in the patient as a preoperative evaluation for AVWS associated with severe mitral regurgitation. A total of 1300 IU (approximately 30 IU/kg) was administered. Plasma samples were collected before the test and at 30 minutes, 6 hours, and 21 hours afterward. VWF multimer analysis was performed and VWF-LMI was evaluated. This series of analyses was approved by the Nara Medical University ethics committee (approval no. 2503).

### Monoclonal anti-ADAMTS13 antibody (HA10)

HA10 was generated using CHO-MK cells^39^ based on the amino acid sequence of HA10^16^. The stable pool was generated at Chitose Laboratory (Kawasaki, Japan), and HA10 was manufactured at Kishi Kasei (Yokohama, Japan).

### *In vitro* PCPS experiment

Two bags, 1/4-inch tubings, and a MERA centrifugal pump (Senko Medical, Tokyo, Japan) were assembled to create a closed system (Fig. 1d). A total of 200 mL of whole blood was obtained from healthy six donors and respectively stored in a CPDA (Citrate-Phosphate-Dextrose-Adenine)- filled bag, which was subsequently introduced into the closed system. HA10 was supplemented at a final concentration of 10 µg/mL when required. During the experiment, pressure head and flow were set to 200 mmHg and 4 L/min, respectively, at a constant speed of 3,000 rpm. Samples (2 mL each) were collected before the perfusion test and at 30, 60, 90, 120, 150, and 180 minutes. Plasma was obtained by centrifugation (2,400 g, 11 min) and stored at −30°C until analysis. A sensitive chromogenic enzyme-linked immunosorbent assay (ELISA) was used to measure ADAMTS13 activity (Kainos Laboratories, Tokyo, Japan). VWF multimer analysis was performed and VWF-LMI was evaluated. This experiment was approved by the Tohoku University ethics committee (approval no. 2024-1-170).

### Plasma-based shear stress assay using an *in vitro* syringe model

An *in vitro* syringe model, which was modified from that described by Hayakawa et al^33^, was assembled by connecting two 1-mL disposable syringes (Henke-Ject® Syringe 1ml Luer Lock (LDS), Tuttlingen, Germany) through a 21-gauge connecting needle with an inner diameter of 0.5 mm (Tsubasa Industry, Tokyo, Japan). A shear stress of 720 dyne/cm^2^ was generated using 442 μL of plasma from various animal spaces (viscosity, 2.0 mPa·s^40^). Every second for 6 minutes (360 times), plasma with or without HA10 (100 μg/mL) was manually transferred back and forth between the syringes.

#### Collection of samples from each animal species

##### Human

The citrated whole blood was obtained from a normal healthy volunteer, which was then centrifuged at 3,000 rpm to obtain plasma. Collection of the sample was approved by the Nara Medical University ethics committee (approval no. 1227).

##### Cynomolgus monkey

The citrated plasma sample from a 3-year-9-month-old female cynomolgus monkey was purchased from HAMRI Co., Ltd. (Koga, Japan). The sample was stored at −80°C until analysis.

##### Porcine

The citrated whole blood obtained from a 3-month-old female domestic porcine was purchased from Ivtech (Kobe, Japan). Platelet-poor plasma (PPP) was prepared by centrifuging whole blood (250 g, 15 min) and the supernatant (2,200 g, 15 min), then stored at −30°C.

##### Bovine and caprine

A 4-week-old male bovine was purchased from Ushizaka (Miyagi, Japan) and a 3-year-old male goat was purchased from Inoue (Gunma, Japan). Those animals were maintained in Dr. Shiraishi’s laboratory. Citrated whole blood was centrifuged (2,400 g, 11 min) to obtain plasma, stored at −30°C. The samples were collected with the approval of the Tohoku University ethics committee (approval no. 2025-AcA-007 (bovine) and 2024-AcA-002 (goat)).

### PCPS-implanted AVWS male monkey experiment

#### PCPS circuit preparation

The PCPS circuit was assembled using a MERA centrifugal pump, a membrane oxygenator (Biocube 2000, Niplo, Osaka, Japan), and 6.0 mm tubings, and was primed with 5% albumin (Japan Blood Products Organization, Tokyo, Japan).

#### Surgical procedure

The experiment was conducted with four male cynomolgus monkeys in total. For induction anesthesia, a mixed solution of ketamine hydrochloride and xylazine was intramuscularly administered. The subject was then intubated and connected to a mechanical ventilator. Maintenance anesthesia was performed throughout the experiment using a mixed gas of oxygen and room air (1:1) with isoflurane vaporized at approximately 1.5-2%. Next, a median sternotomy was made and the ascending aorta was exposed. After administration of heparin (300 U/kg), a partial clamp was applied to the ascending aorta. An arterial return cannula (3.5 mm) was then inserted end-to-side on the aorta using prolene 5-0. Next, a venous drainage cannula (4.0 mm) was inserted into the right atrium through the apical incision and secured using prolene 5-0. The both cannulas were then connected to the circuit. HA10 with a final concentration of 10 μg/mL was administered to two animals, whereas nothing was given to the remaining two. The perfusion experiment was initiated at an initial pump speed of 3,000 rpm. Blood samples (4 mL each) were collected before cannulation, immediately before perfusion (set as baseline), and at 30, 60, 120, and 180 minutes. Citrated blood was centrifuged (3,000 rpm, 10 min, 4°C) and the plasma was store at −80°C until analysis. The study was approved by the Institutional Animal Care and Use Committees of Nissei Bilis (approval no. 2023-182).

### Single dose of HA10 in cynomolgus monkeys

#### Preliminary non-GLP single-dose study

HA10 was intravenously administered at 6.5 or 32.5 mg/kg to two male cynomolgus monkeys per group, targeting plasma concentrations of 100 and 500 μg/mL. Hematological and biochemical parameters were evaluated at 1, 3, and 6 days post-administration. Histopathological examinations of the heart and brain were conducted on day 14. Blood sampling for toxicokinetic studies was performed at 0.167, 1, 2, 4, 6, 8 hours and at 1, 3, 6 days after HA10 administration.

#### Definitive single-dose GLP study

HA10 was intravenously administered at 3 and 25 mg/kg to three males per group. The tested doses of the GLP study were set based on the preliminary study results. The 3 mg/kg dose was selected to sustain plasma levels above the estimated effective concentration (10 μg/mL) for approximately 1 week. The control group (n = 3) received 146 mM purified sucrose/0.05% (w/v) polysorbate 80 in PBS (-), administered identically to HA10. During the observation period, blood biochemical analyses were performed on days −8 (as baseline), 1, 3, 6, and 14; necropsy and histopathological examinations were conducted on day 14. Blood sampling for toxicokinetic studies was performed at 0.167, 1, 2, 4, 6, 8 hours and at 1, 3, 6, 14 days after HA10 administration.

The toxicity studies were approved by the Institutional Animal Care and Use Committees of Shin Nippon Biomedical Laboratories (approval no. IACUC722-045). The facility is accredited by AAALAC international.

### ADAMTS13 inhibition assay by ELISA

ADAMTS13 activity was measured by the aforementioned ELISA. The concentration-dependent inhibitory effect of HA10 on ADAMTS13 activity was determined by incubating standard human plasma (Siemens Healthineers) for an hour with a final HA10 concentration of 0.05-100 µg/mL. The values at dose 0 control was also measured to calculate the inhibitory rate. The IC_50_ value was estimated using R software (version 4.3.2) packages.

### Statistical analysis

In the *in vitro* PCPS experiment, the equality of variances was first assessed using the F-test, followed by the student’s t-test to compare means between samples with and without HA10. In the definitive HA10 single-dose GLP study, the homogeneity of variances was evaluated using Bartlett’s test. If variances were equal, multiple comparisons between the control group and each test group were performed using Dunnett’s test; otherwise, Miller’s test was used. p < 0.05 was considered significant. Statistical analyses for general blood tests in the GLP study were performed using MiTOX (Mitsui E&S Systems Research Inc., Chiba, Japan), while all other analyses were conducted using EZR^41^.

### Expression and purification of ADAMTS13D in tabacco BY-2 cells

Recombinant ADAMTS13D was produced in *Nicotiana tabacum* BY-2 cells using an estradiol-inducible tobamovirus vector^42–44^. A codon-optimized ADAMTS13D sequence (residues X– X) was substituted for ORF8 between the PmeI and BstEII sites of pBICLBSER-ToMV-SP-His-ORF8, generating pBICLBSER-ToMV-SP-His-ADAMTS13D-HEDL. The construct encoded an N-terminal signal peptide and 8×His tag and a C-terminal HEDL ER-retention signal.

The construct was introduced into BY-2 cells stably expressing XVE. Expression was induced with 10 μM 17β-estradiol 2 days after subculture, and cells were harvested 4 days later. Freeze–thawed cells were sonicated in 6 M guanidine hydrochloride, 20 mM sodium phosphate, 500 mM NaCl, and 20 mM imidazole (pH 7.4), followed by centrifugation at 64,000 × g for 10 min at 4°C.

The supernatant was applied to a His GraviTrap column (Cytiva), and protein was eluted with 4 M urea, 20 mM sodium phosphate, 500 mM NaCl, and 500 mM imidazole (pH 7.4). The eluate was further purified by size-exclusion chromatography in the same buffer and refolded by stepwise dialysis against 0.02× PBS.

### Expression and purification of ADAMTS13D and VWF73 in *E. coli*

Expression plasmids for a N-terminal His-tagged ADAMTS13D and mutants were transformed into *E. coli* BL21(DE3) cells. *E. coli* cells were cultured in LB/Amp medium at 37°C, and proteins were expressed with 0.5 mM isopropyl-β-D-thiogalactopyranoside (IPTG) at 20°C overnight. For ^15^N-labeling or ^13^C,^15^N-double labeling, *E. coli* cells were respectively cultured in M9/Amp medium supplemented with ^15^NH_4_Cl, or ^15^NH_4_Cl, ^13^C-glucose and ISOGRO-^13^C,^15^N Powder. For the preparation of 75% deuterated (75d)-ADAMTS13D and 100% deuterated (100d)-ADAMTS13D, *E. coli* cells were cultured in M9/Amp medium containing 75% or 100% D_2_O, respectively, supplemented with either a glucose mixture composed of 75% (w/w) D-glucose (1,2,3,4,5,6,6-D7) and 25% (w/w) unlabeled D-glucose for 75d-ADAMTS13D or 100% (w/w) D-glucose (1,2,3,4,5,6,6-D7) for 100d-ADAMTS13D. The labeling proteins were expressed with 0.5 mM IPTG at 37°C overnight. After protein expression, *E. coli* cells were collected, suspended in lysis buffer (50 mM Tris-HCl, pH 7.5, 500 mM NaCl, 1% Triton X-100, and protease inhibitor) with 0.2 mg/mL lysozyme, sonicated, and centrifuged at 12,000×g for 15 min at 4°C. Inclusion bodies were washed with lysis buffer and collected by centrifugation at 12,000×g for 15 min at 4°C. The refolding of inclusion bodies was performed according to the previous report^45^ as follows: inclusion bodies were solubilized in solubilization buffer (100 mM NaH_2_PO_4_, 10 mM Tris-HCl, pH 8.0, and 8M urea), diluted 20-fold in refolding buffer (50 mM MES, pH 6.0, 240 mM NaCl, 10 mM KCl, 1 mM EDTA, 0.5 M Arginine, 0.4 M Sucrose, 0.5% Triton X-100, 0.05% PEG-20,000, 1 mM reduced glutathione, and 0.1 mM oxidized glutathione), and stirred at room temperature for 72 h. After dialysis against dialysis buffer (20 mM Tris-HCl, pH 8.0, and 150 mM NaCl), recombinant proteins were purified using Ni-NTA agarose (FUJIFILM Wako Pure Chemical Corporation, Osaka, Japan) with wash buffer (PBS) and elution buffer (20 mM Tris-HCl, pH 8.0, 200 mM NaCl, and 250 mM imidazole), followed by gel filtration using Superdex 200 pg with PBS.

The GST-VWF73-His expression plasmid was transformed into *E. coli* BL21(DE3) cells. *E. coli* cells were cultured in LB/Amp medium at 37°C, and proteins were expressed with 0.5 mM IPTG at 30°C for 4 h. For ^15^N-labeling or ^13^C,^15^N-double labeling, *E. coli* cells were respectively cultured in M9/Amp medium supplemented with ^15^NH_4_Cl, or ^15^NH_4_Cl, ^13^C-glucose and ISOGRO-^13^C,^15^N Powder. Cells were suspended in lysis buffer (20 mM Tris-HCl, pH 7.5, 150 mM NaCl, and protease inhibitor), sonicated, and centrifuged at 12,000×g for 30 min at 4°C. The supernatant was applied to Glutathione Sepharose 4B beads (Cytiva, Tokyo, Japan) and subsequently purified using a Ni-NTA agarose column. The GST-tag was cleaved with PreScission Protease (Cytiva, Tokyo, Japan) at 4°C for 16 h. The cleaved sample was incubated with Glutathione Sepharose 4B beads to remove the GST-tag and protease, and further purified using Superdex 200 pg with PBS.

### Verification of the disulfide bond pattern

The disulfide linkage pattern of ADAMTS13D was determined according to a previously described method^46^. In brief, ADAMTS13D was diluted to 0.5 mg/mL in 20 mM Tris-HCl (pH 8.0) and subjected to a two-step digestion with trypsin (1:100, 37°C overnight) followed by Asp-N (1:50, 37°C overnight) or ProAlanase (1:100, 37°C for 2 h) under nonreducing conditions. The digested samples were acidified with 1% formic acid, desalted using GL-Tip SDB and GL-Tip GC (GL Sciences Inc.), dried under vacuum, and analyzed by MALDI-TOF-MS (UltrafleXtreme, Bruker Daltonics, Billerica, MA, USA). Data were analyzed using BioTools 3.2 (Bruker Daltonics, Billerica, MA, USA).

### mA10-binding assays

Equal molar amounts of ADAMTS13D and mA10 (1.2 μM each) were mixed in PBS. After incubation with Dynabeads Protein G at 4°C for 30 min, the beads were washed three times with PBS and analyzed by SDS-PAGE.

To evaluate ADAMTS13D mutants, equal molar amounts of wild-type and mutant proteins (1.8 μM each) were subjected to SDS-PAGE and transferred to polyvinylidene difluoride membranes. Membranes were blocked with TBS-T (20 mM Tris-HCl pH 7.4, 150 mM NaCl, and 0.05% Tween 20) containing 5% skim milk and incubated with mA10 (5 μg/mL). After washing with TBS-T buffer, membranes were incubated with anti-mouse IgG HRP-linked antibody (1:10000, RGAM001, lot 20001141, Proteintech, Rosemont, IL, USA). Blots were visualized using Clarity Western ECL substrate (Bio-Rad Laboratories, Hercules, CA, USA) and images were acquired using the FUSION system (Vilber Bio Imaging, Marne-la-Vallée, France).

### NMR experiments

Backbone resonance assignment experiments were performed using NMR spectra measured on a Bruker AVANCE NEO 800 MHz spectrometer equipped with a CPTCI probe. ^13^C,^15^N-labeled ADAMTS13D or VWF73 at a concentration of 0.4 mM dissolved in PBS and 5% ^2^H_2_O were used. The sample temperature was set at 25°C for both samples. The backbone resonance assignments were carried out using the following triple resonance spectra: HNCO, HNCA, HN(CO)CA, HNCACB, CBCA(CO)NH, C(CO)NH, and ^15^N-edited NOESY. The data were processed using Topspin and analyzed using Sparky^47^.

For the titration experiments with ADAMTS13D, the NMR spectra were measured using a Bruker AVANCE 500 MHz spectrometer equipped with a BBFO probe. ^15^N-labeled ADAMTS13D at a concentration of 0.1 mM dissolved in PBS and 5% ^2^H_2_O was used. The sample temperature was set at 25°C. Backbone ^1^H-^15^N correlation spectra were obtained *via* TROSY-HSQC experiments. HA10 was titrated into ADAMTS13D at molar ratios of 0.1, 0.2, 0.5, and 1. For the titration experiments with VWF73, the NMR spectra were measured on the Bruker AVANCE 500 MHz spectrometer equipped with the BBFO probe. ^15^N-labeled VWF at a concentration of 0.1 mM dissolved in PBS and 5% ^2^H_2_O was used. The sample temperature was set at 25°C. Backbone ^1^H-^15^N correlation spectra were obtained *via* SOFAST-HMQC experiments. HA10 was titrated into VWF at molar ratios of 0.2, 0.5, and 1 in the absence of ADAMTS13D, and at ratios of 0.5, 1, and 2 in the presence of 0.1 mM ADAMTS13D. The data were processed using NMRPipe^48^ and analyzed using POKY^49^. The error bar for the intensity ratio (ΔR) was estimated from the signal-to-noise ratios using the equation: ΔR = R(SNR_ref_^-2^ + SNR_tit_^-2^)^1/2^ where R is the intensity ratio, and SNR and SNR are signal-to-noise ratios of the peaks on the reference and titrated spectrum, respectively^50^.

### Molecular modeling of ADAMTS13-HA10 complex

Docking simulations, including information from NMR and mutagenesis experiments, were performed using HADDOCK 2.4 web server^51^. The atomic coordinate of X-ray crystal structure of ADAMTS13 (PDB 6QIG) was used for the initial model. The HA10 model was built with sequences from variable regions connected by 10-repeats of the GS linker, and the coordinates were predicted using AlphaFold2^52^ *via* LocalColabFold^53^. Active residues for ADAMTS13 were defined as (310, 332, 338, 339, 349, and 371), which were significantly perturbed in the NMR experiment. The additional active residues were defined as (308, 309, 340, 350, and 352) based on the mutagenesis experiments. Passive residues for ADAMTS13 were automatically defined around the active residues. Active residues for HA10 were not defined. Passive residues for HA10 were manually defined at residues in the complementarity-determining region^16^. Buried residues (with a relative solvent accessibility smaller than 15%) were removed from the selection of the active and passive residues for ADAMTS13 and HA10. The 162 of 200 calculated structures were clustered on the basis of the algorithm implemented in the HADDOCK server. The representative structure was selected from the best cluster with the lowest HADDOCK score (−123.1 +/− 5.3), the highest cluster size (71) and the most negative z score (−2.2).

### Analytical ultracentrifugation (AUC)

All samples for AUC measurements were prepared in PBS. Solutions containing full-length A10 (mA10 or HA10) alone or ADAMTS13D alone were prepared at 1.5 mg/mL. Mixtures of full-length A10 and ADAMTS13D were prepared at molar ratios of 1:0.5, 1:1, 1:2, and 1:3, with the concentration of full-length A10 fixed at 1.5 mg/mL. The mA10–ADAMTS13D series was measured in H_2_O-based PBS, whereas the HA10–ADAMTS13D series was measured in PBS prepared with 100% D_2_O.

AUC measurements were performed with a ProteomeLab XL-I (Beckman Coulter, USA). The samples were loaded into a cell with a 12 mm optical-pathlength and were measured using Rayleigh interference optics at a rotor speed of 45,000 rpm and temperature of 25°C. The time evolution of the sedimentation data was analyzed with Lamm formula using SEDFIT software (version 15.01c)^54^. The weight concentration distribution *c*(*s*_20,w_) was obtained as a function of the sedimentation coefficient which was normalized to the value at 20 °C in pure H_2_O *s*_20,w_.

### Small-angle X-ray scattering (SAXS)

All samples for SAXS measurements were prepared in PBS. Solutions containing full-length A10 alone (mA10 or HA10) or ADAMTS13D alone were prepared at 1.5 mg/mL. Mixtures of full-length A10 and ADAMTS13D were prepared at a molar ratio of 1:2, with the concentration of full-length A10 at 1.5 mg/mL. The mA10–ADAMTS13D series was measured in H_2_O-based PBS, whereas the HA10–ADAMTS13D series was measured in PBS prepared with 100% D_2_O.

SAXS measurements were performed using a laboratory-based instrument NANOPIX (RIGAKU, Japan) equipped with a high-brilliance point-focused generator of a Cu-Kα source MicroMAX-007 HFMR (Rigaku, Japan) (wavelength = 1.54 Å). The sample-to-detector distances were set to be 1330 and 300 mm, with which the covered *q*-range was 0.01 Å^-1^ ≤ *q* ≤ 0.50 Å^-1^ (*q*: magnitude of scattering vector). All measurements were performed at 25°C. Two-dimensional scattering patterns were converted to one-dimensional scattering profiles using SAngler software (version 2.1.71) ^55^.

AUC-SAXS treatment was applied to eliminate the influence of aggregates from experimental SAXS profile. Details of the AUC-SAXS treatment have been described in existing studies^56,57^.

### Small-angle neutron scattering (SANS)

To selectively characterize the solution structure of full-length A10 (mA10 or HA10) within the A10–ADAMTS13D complex, hydrogenated (h)-A10 and 75% deuterated (75d)-ADAMTS13D were prepared in PBS containing 100% D_2_O. Under these conditions, (h)-A10 contributes to the neutron scattering signal, whereas (75d)-ADAMTS13D is contrast-matched to the solvent and is therefore effectively invisible. This approach is referred to as inverse contrast matching^58^. Mixtures of (h)-A10 and (75d)-ADAMTS13D were prepared at a molar ratio of 1:2, with the concentration of (h)- A10 at 1.5 mg/mL. As a control sample, a solution containing (h)-A10 alone was prepared at 1.5 mg/mL.

To selectively characterize the solution structure of ADAMTS13D within the Fab of HA10– ADAMTS13D complex, hydrogenated (h)-Fab and 100% deuterated (100d)-ADAMTS13D were prepared in PBS containing 42% D_2_O. Under these conditions, (100d)-ADAMTS13D contributes to the neutron scattering signal, whereas (h)-Fab is contrast-matched to the solvent and is therefore effectively invisible. This approach is referred to as contrast matching^59^. Mix solutions of (h)-Fab and (100d)-ADAMTS13D were prepared to be 1:1 of molar ratio, where the concentration of (100d)- ADAMTS13D was 0.22 mg/mL. As a control sample, (100d)-ADAMTS13D solo solution was prepared at 2.3 mg/mL. Deuteration of ADAMTS13D was conducted with the manner developed in the previous study^60^.

SANS measurements were performed with SANS-U at JRR-3 (Institute for Solid State Physics, The University of Tokyo and the Japan Atomic Energy Agency) for mA10–ADAMTS13D series and D22 and D33 at Institut Laue-Langevin for HA10–ADAMTS13D series. A neutron beam with a wavelength (λ) of 6.0 Å and its distribution (Δλ/λ) of 10% was irradiated to the samples at all instruments. The sample-to-detector distances were set to be 8000, 4000, and 1030 mm at SANS-U (magnitude of scattering vector *q* range of 0.007 – 0.3 Å^-1^), 5600 and 1400 mm at D22 (*q* range of 0.008 – 0.4 Å^-1^), and 4000 and 1400 mm at D33 (*q* range of 0.01 – 0.4 Å^-1^). All measurements were performed at 25°C. Two-dimensional scattering patterns were converted to one-dimensional scattering profiles using GRASP software (version 11.05c).

### Structural modeling based on SAXS

As a structural model of full-length HA10, a homology model was generated using SWISS-MODEL^61^, with the crystal structure of an antibody belonging to a subclass similar to that of A10 (PDB ID: 1IGT) as the template. Referring to the interaction interface between ADAMTS13D and A10 domain identified by NMR and docking simulation analysis, we constructed a full-length docking model. Finally, normal mode analysis optimization (NMA optimization) was performed for the full-length docking model using the Pepsi-SAXS-NMA software (version 3.0)^62^, yielding a three-dimensional model that successfully reproduces the SAXS scattering profile of the full length A10-ADAMTS13D complex at a 1:2 binding stoichiometry.

The agreement between the calculated scattering profile *I*_cal_(*q*) from model structures and the experimental one *I*_exp_(*q*) was evaluated using the *χ*^2^-value in the following.

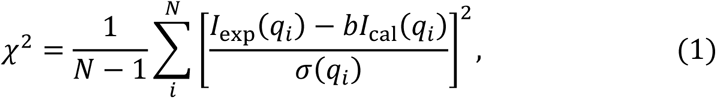

where *N* is the number of points in the scattering profile, *σ*(*q*) are the experimental errors, and *b* is the scaling factor.

### Analysis of the redox states of ADAMTS13 disintegrin-like domain

Recombinant ADAMTS13 disintegrin-like domain (100 μM) was incubated with reduced glutathione (GSH; 0–6 mM) in PBS at 30 °C. At the indicated time points, free thiols were alkylated by addition of 1 mM iodoacetamide (FUJIFILM Wako Pure Chemical Corporation) to quench further thiol–disulfide exchange reactions.

For the counter alkylation analysis by mass spectrometry, the samples after iodoacetamide treatment were subjected to TCA precipitation to remove excess iodoacetamide. The precipitated proteins were dissolved in 50mM ammonium bicarbonate and 1% SDS solution, reduced by TCEP (to a final concentration of 2.5mM), and methylthiolated by the *S*-Methyl methanethiosulfonate (to a final concentration of 20mM). Then, proteins were digested using the SP3 method^63^. After the digestion, peptides were desalted by GL-Tip SDB (GL sciences, Japan), and measured by nanoLC-ESI-mass-spectrometer system (Easy-nLC1000 nanoflow liquid chromatograpy system and Q-Exactive tandem mass spectrometer) in a data-dependent acquisition mode. Annotation of methylthiolated peptides was performed using the FragPipe platform (ver. 22) bundled with MSFragger search engine^64^, and quantification was performed using Skyline (ver. 24)^65^.

### Accessible surface area analysis

The accessible surface area (ASA) of cysteine sulfur atoms in the ADAMTS13 disintegrin-like domain was calculated from the crystal structure of ADAMTS13 (PDB ID: 6QIG) using the GETAREA program^66^. ASA values were calculated for each cysteine sulfur atom involved in disulfide bond formation and mapped onto the ADAMTS13 disintegrin-like domain structure for visualization.

### Isothermal titration calorimetry

ITC experiments were performed using an iTC200 calorimeter (Malvern Panalytical) in PBS. ADAMTS13D (100 μM) was titrated into the sample cell containing HA10 (10 μM) or GST-VWF73-His (20 μM). Measurements were performed at 25 °C with preliminary 0.4 μL injection followed by 19 subsequent 2 μL additions. Data were analyzed using MicroCal PEAQ-ITC Analysis Software.

## Acknowledgements

We thank Takeshi Yokoyama (Tohoku University), Yuta Aizawa (Tohoku University), Yoshikazu Tanaka (Tohoku University), Koji Yonekura (Tohoku University), Yu Hirano (QST), and Taro Tamada (QST) for their technical support in structural analyses. The experiment using an *in vitro* syringe model was performed with valuable advice from Dr. Koichi Kokame (National Cerebral and Cardiovascular Center, Suita, Japan). We also thank Ayumi Kanemaru, Masanori Fujiwara, and Izumi Kamo at CRIETO (Clinical Research Innovation and Education Center Tohoku University Hospital), Tohoku University for their administrative support in project management and regulatory interviews, the treating physicians in the Department of Pediatrics at Nara Medical University for kindly providing the patient samples, Justin Gordin (UT Southwestern Medical Center) for his valuable suggestions, and Keren-Happuch E for her critical reading of the manuscript.

SANS measurements were conducted with SANS-U at JRR-3 (Institute for Solid State Physics, The University of Tokyo and the Japan Atomic Energy Agency) under proposal No. 24404, and D22 and D33 at Institut Laue-Langevin under proposal No. 8-03-1137. AUC and SAXS measurements were conducted at KURNS under proposal Nos. R4129, R7054, and R8059. This study was partially supported by Platform Project for Supporting Drug Discovery and Life Science Research [Basis for Supporting Innovative Drug Discovery and Life Science Research (BINDS)] from AMED (JP24ama121001j0001 to M.S.), and by AMED under Grant Number JP17ek0109246 and JP22ek0109579 (to M. Matsumoto).

## Contributions

M. Matsumoto and E.M. conceptualized the study. M. Matsumoto, E.M., K. Saito, N.I. and Y.S. designed the experiments. T.U. and S.H. treated the patient with Impella and provided the patient samples. Y.S. performed *in vitro* PCPS experiment and surgical procedures in the PCPS-implanted AVWS male monkey experiment. R.I., M.K. and K.N. performed the PCPS-implanted AVWS male monkey experiment. T.Y. contributed to the experimental design and provided methodological guidance in the in vitro PCPS experiment and the PCPS-implanted AVWS male monkey experiment. Y.H., H.K. and T.S. performed NMR experiments. N.I., R.S., M. Mori, T.I., M.N., M.T. and N.T. prepared recombinant proteins. M. Oda performed ITC experiments. S.K., M. Okumura and T.N. performed redox analyses. A.M., L.P., K.M., A.O. and M.S. performed AUC, SAXS, and SANS experiments. N.I. verified the disulfide bond pattern. N.I., M.N., K. Sakai and A.H. performed mA10-binding assays. K. Saito, N.I., Y.S. and E.M. drafted and revised the manuscript. All authors read and approved the final manuscript.

