## Supplemental figure 1-9 for "Calibrated ADAMTS13 inhibition prevents cardiovascular shear coagulopathy"

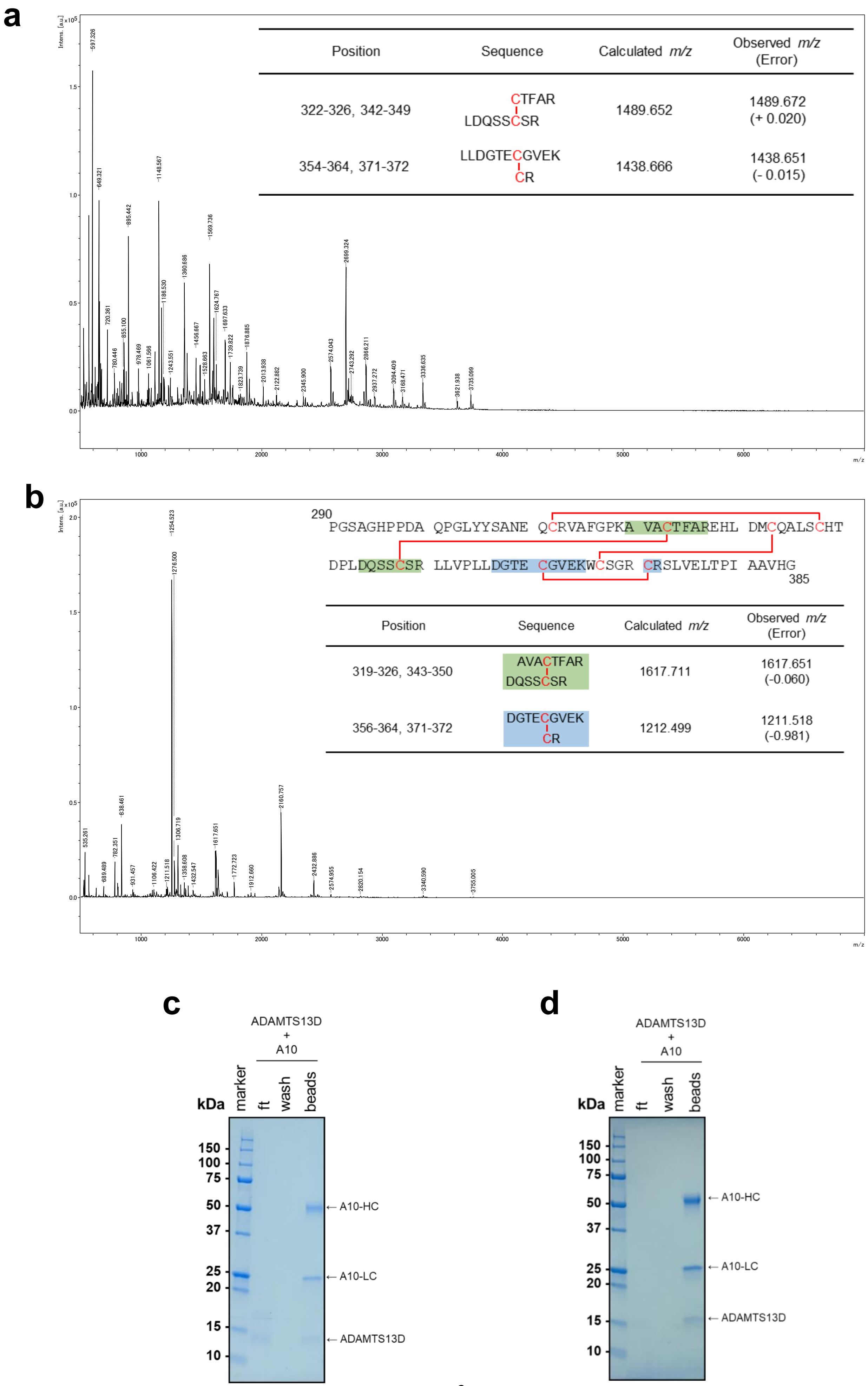
a
Position | Sequence | Calculated m/z | Observed m/z (Error)
322-326, 342-349 | CTFAR / LDQSSCSR | 1489.652 | 1489.672 (+ 0.020)
354-364, 371-372 | LLDGTECGVEK / CR | 1438.666 | 1438.651 (- 0.015)
b
290
PGSAGHPPDA QPGLYYSANE QCRVAFGPKA VACTFAREHL DMCQALSCHT
DPLDQSSCSR LLVPLLDGTE CGVEKWCSGR CRSLVELTPI AAVHG
385
Position | Sequence | Calculated m/z | Observed m/z (Error)
319-326, 343-350 | AVACTFAR / DQSSCSR | 1617.711 | 1617.651 (-0.060)
356-364, 371-372 | DGTECGVEK / CR | 1212.499 | 1211.518 (-0.981)
c
ADAMTS13D + A10
kDa
marker
ft
wash
beads
150
100
75
50
37
25
20
15
10
A10-HC
A10-LC
ADAMTS13D
d
ADAMTS13D + A10
kDa
marker
ft
wash
beads
150
100
75
50
37
25
20
15
10
A10-HC
A10-LC
ADAMTS13D

**Extended Data Fig. 1: Characterization of ADAMTS13D prepared using BY-2 cells and *E. coli* expression systems. a,b,** Mass spectrometric analysis of disulfide bond connectivity in ADAMTS13D prepared using BY-2 cells (**a**) and *E. coli* (**b**). **c**,**d**, Binding of ADAMTS13D prepared using BY-2 cells (**c**) and *E. coli* (**d**) to mA10.

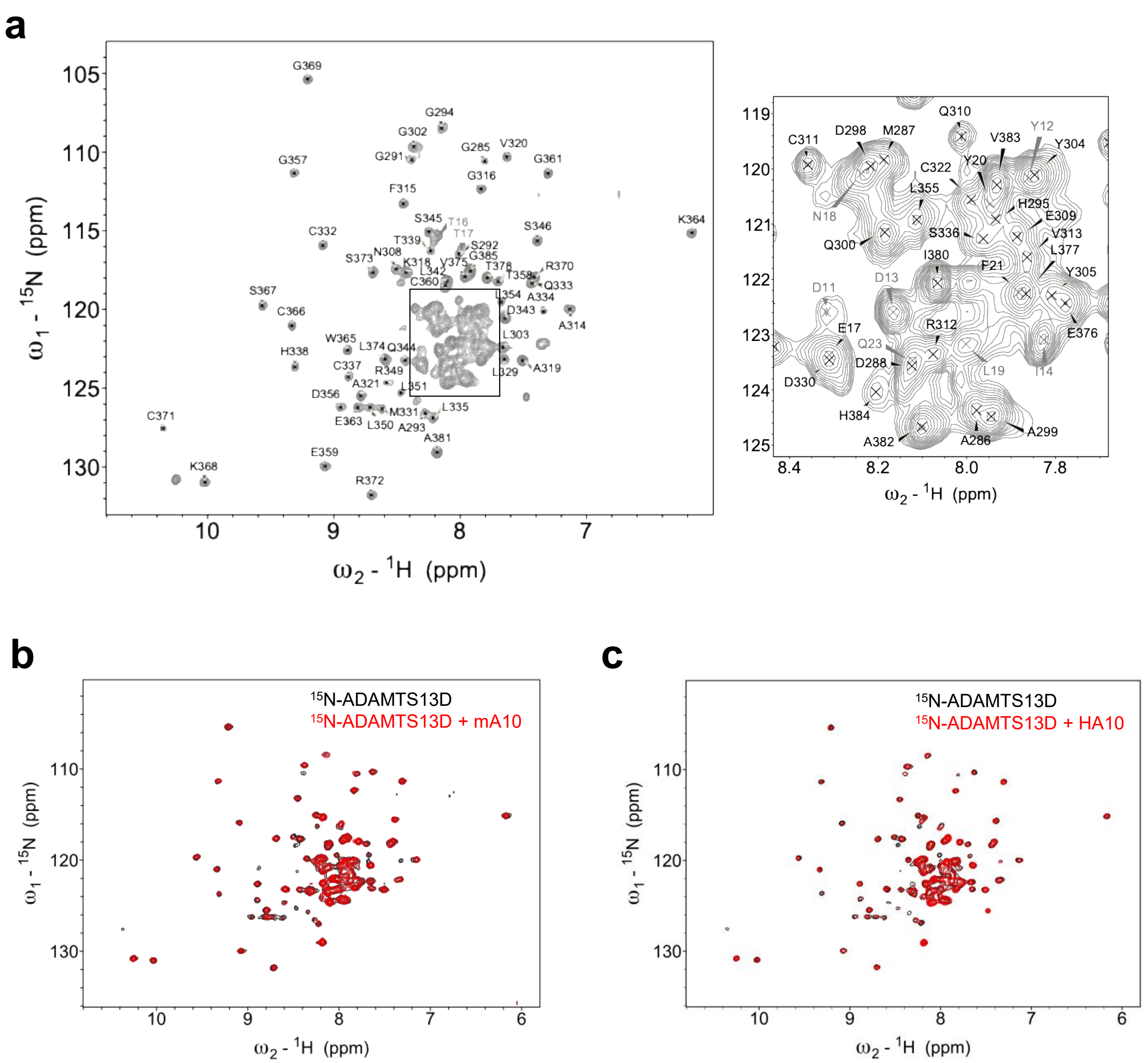


**Extended Data Fig. 2: NMR analysis of the interactions of stable isotope-labeled ADAMTS13D with mA10 and HA10. a**, NMR signal assignments of ADAMTS13D. **b,c**, Overlay of $^{1}H$-$^{15}N$ NMR spectra of $^{15}N$-labeled ADAMTS13D in the absence and presence of mA10 (**b**) or HA10 (**c**).

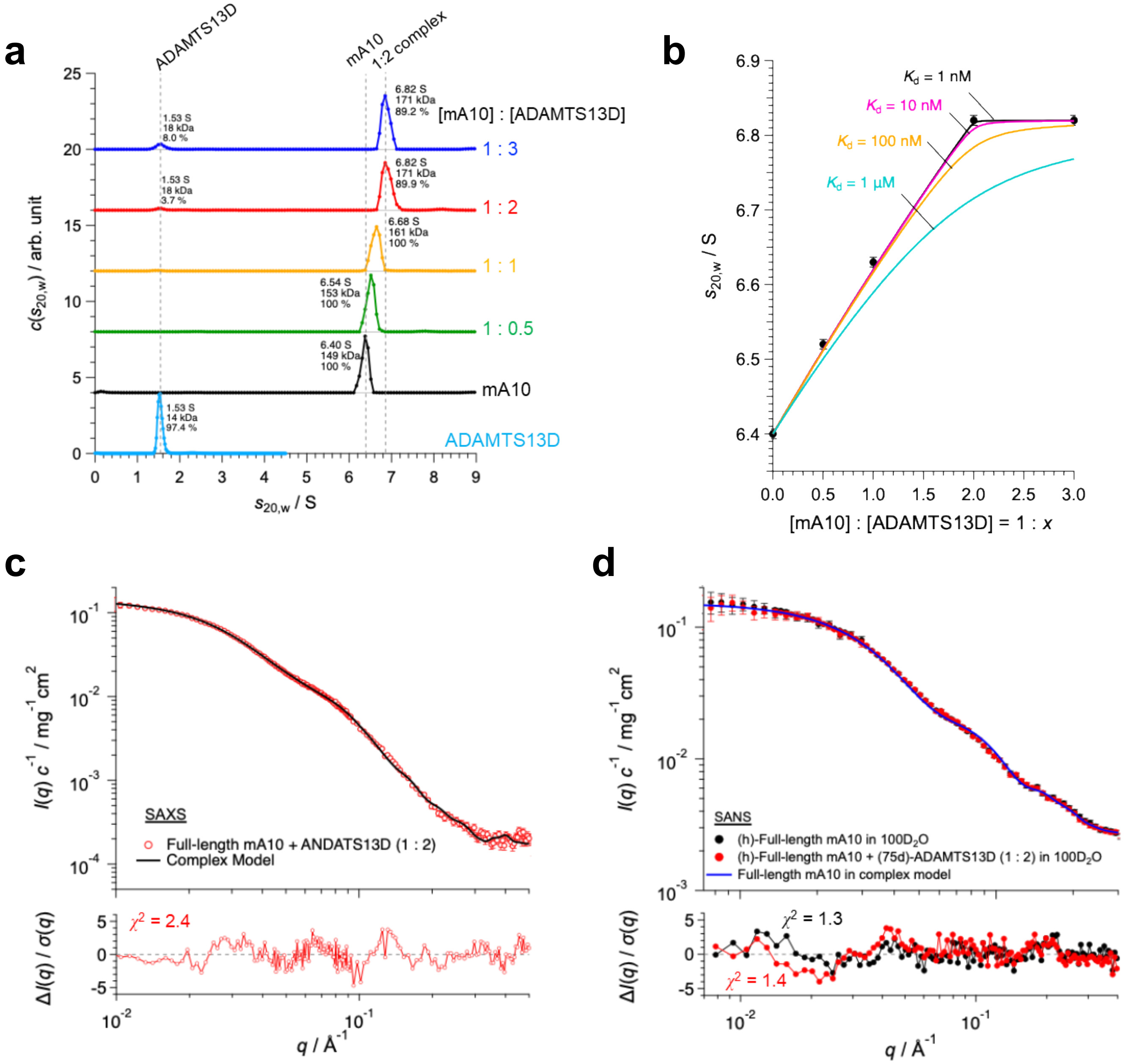

a
ADAMTS13D
mA10
1:2 complex
[mA10] : [ADAMTS13D]
6.82 S
171 kDa
89.2 %
1.53 S
18 kDa
8.0 %
1 : 3
6.82 S
171 kDa
89.9 %
1.53 S
18 kDa
3.7 %
1 : 2
6.68 S
161 kDa
100 %
1 : 1
6.54 S
153 kDa
100 %
1 : 0.5
6.40 S
149 kDa
100 %
mA10
1.53 S
14 kDa
97.4 %
ADAMTS13D
$c(s_{20,w})$ / arb. unit
$s_{20,w}$ / S
b
$K_d$ = 1 nM
$K_d$ = 10 nM
$K_d$ = 100 nM
$K_d$ = 1 μM
$s_{20,w}$ / S
[mA10] : [ADAMTS13D] = 1 : $x$
c
SAXS
Full-length mA10 + ANDATS13D (1 : 2)
Complex Model
$I(q)\ c^{-1}$ / mg$^{-1}$ cm$^{2}$
$\chi^2$ = 2.4
$\Delta I(q) / \sigma(q)$
$q$ / Å$^{-1}$
d
SANS
(h)-Full-length mA10 in 100D$_2$O
(h)-Full-length mA10 + (75d)-ADAMTS13D (1 : 2) in 100D$_2$O
Full-length mA10 in complex model
$I(q)\ c^{-1}$ / mg$^{-1}$ cm$^{2}$
$\chi^2$ = 1.3
$\chi^2$ = 1.4
$\Delta I(q) / \sigma(q)$
$q$ / Å$^{-1}$

**Extended Data Fig. 3. Characterization of a full-length mA10-ADAMTS13D complex in solution and three-dimensional modeling. a.** The weight concentration distribution $c(s_{20,w})$ as a function of sedimentation coefficient $s_{20,w}$ obtained by analytical ultracentrifugation (AUC) for ADAMTS13D, full-length mA10, and mix solutions of full-length mA10 + ADAMTS13D with various mixing molar ratios. **b.** Mixing molar ratio dependence of the $s_{20,w}$-value corresponding to full-length mA10 and full-length mA10-ADAMTS13D complex. Peak $s_{20,w}$-values were determined by Gaussian fitting of $c(s_{20,w})$, and the error bars represent the standard errors of the fitted peak positions. The solid lines represent the mixing-ratio dependence of the weight-average sedimentation coefficient for full-length mA10 and the full-length mA10–ADAMTS13D complex, calculated using the method of the Dam and Schuck method[36]. The black, magenta, yellow, and cyan lines correspond to calculated weight-average sedimentation coefficients for dissociation constant $K_d$ values of 1 nM, 10 nM, 100 nM, and 1 μM, respectively. Because the calculated curves become relatively insensitive to $K_d$ value in the submicromolar range, the $K_d$ value could not be determined precisely from the AUC data alone. Nevertheless, the experimental weight-average sedimentation coefficients were better reproduced by the curves calculated for submicromolar $K_d$ values, supporting a relatively high-affinity interaction. At mixing molar ratios above 1:2, the sedimentation coefficient saturated at 6.82 S, corresponding to a complex with a binding stoichiometry of 1:2. **c.** Red circles show the experimental SAXS profile for a full-length mA10-ADAMTS13D complex with a binding stoichiometry of 1:2. Black line represents the calculated scattering profile for the full-length mA10-ADAMTS13D complex model. The scattering intensity is normalized by the concentration. The residuals between the experimental and calculated scattering profiles are shown in the lower panels. **d.** Partial structural analysis of the full-length mA10 in full-length mA10-ADAMTS13D complex using inversed contrast matching SANS. Black and red circles represent the scattering profile of hydrogenated (h)-full-length mA10 in 100%$D_2O$ and mix solution of (h)-full-length mA10 and 75% deuterated (75d)-ADAMTS13D in 100%$D_2O$, respectively. The scattering intensities are normalized by concentration of (h)-full-length mA10 ($c_{HA10}$). Because the scattering length density of (75d)-ADAMTS13D matches that of 100% $D_2O$, (75d)-ADAMTS13D is invisible to neutron scattering. The agreement between the two scattering profiles means that the partial structure of full-length mA10 in the complex is identical to full-length mA10 alone in solution. Blue solid line represents the calculated scattering profile for full-length mA10 in mA10-ADAMTS13D complex. The residuals between the experimental and calculated scattering profiles are shown in the lower panels.

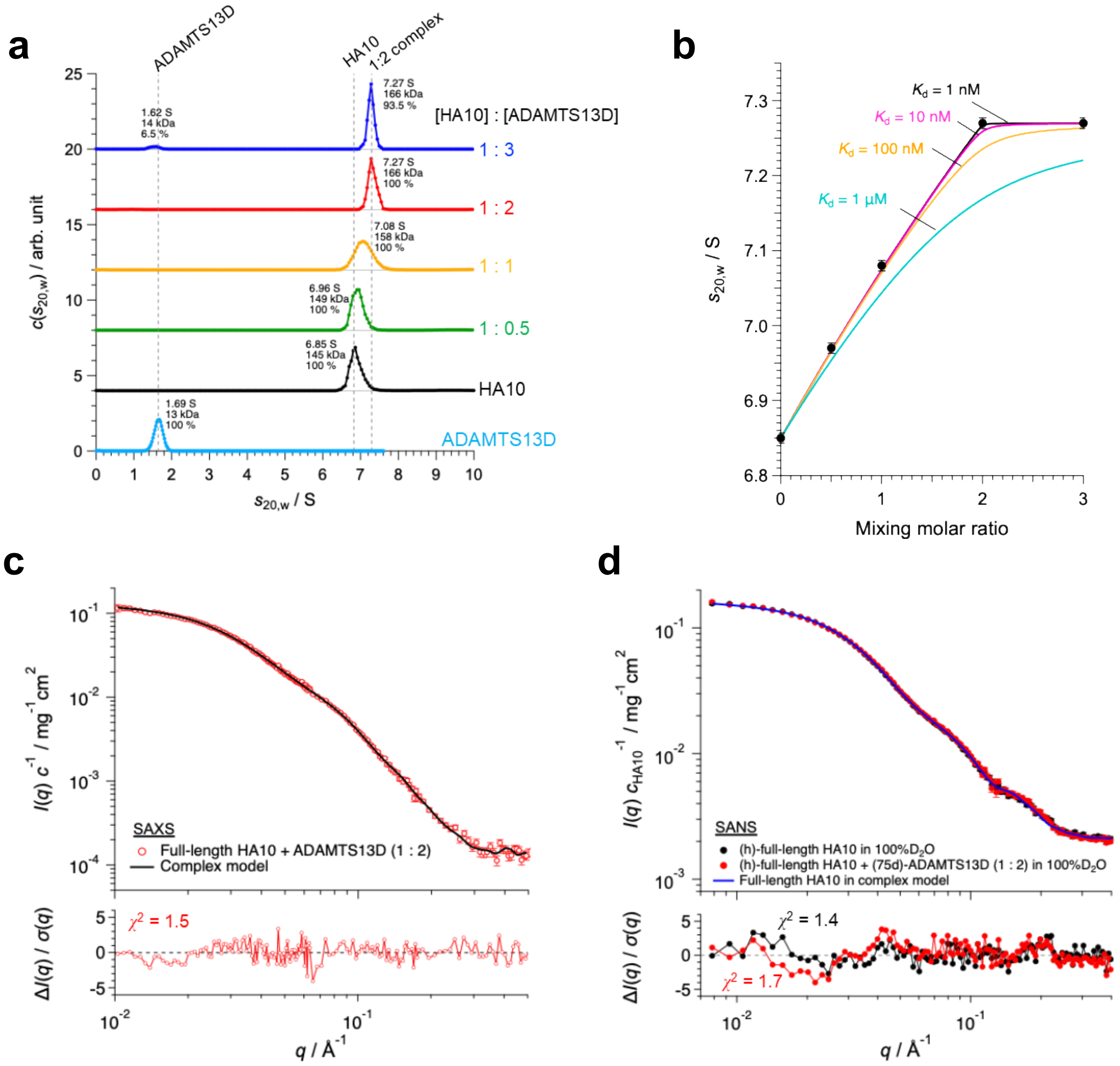

a
ADAMTS13D
HA10
1:2 complex
[HA10] : [ADAMTS13D]
1.62 S
14 kDa
6.5 %
7.27 S
166 kDa
93.5 %
1 : 3
7.27 S
166 kDa
100 %
1 : 2
7.08 S
158 kDa
100 %
1 : 1
6.96 S
149 kDa
100 %
1 : 0.5
6.85 S
145 kDa
100 %
HA10
1.69 S
13 kDa
100 %
ADAMTS13D
$c(s_{20,w})$ / arb. unit
$s_{20,w}$ / S
b
$K_d$ = 1 nM
$K_d$ = 10 nM
$K_d$ = 100 nM
$K_d$ = 1 μM
$s_{20,w}$ / S
Mixing molar ratio
c
SAXS
Full-length HA10 + ADAMTS13D (1 : 2)
Complex model
$I(q)\ c^{-1}$ / mg$^{-1}$cm$^2$
$\chi^2$ = 1.5
$\Delta I(q) / \sigma(q)$
$q$ / Å$^{-1}$
d
SANS
(h)-full-length HA10 in 100%D$_2$O
(h)-full-length HA10 + (75d)-ADAMTS13D (1 : 2) in 100%D$_2$O
Full-length HA10 in complex model
$I(q)\ c_{HA10}^{-1}$ / mg$^{-1}$cm$^2$
$\chi^2$ = 1.4
$\chi^2$ = 1.7
$\Delta I(q) / \sigma(q)$
$q$ / Å$^{-1}$

**Extened Data Fig. 4: Characterization of a full-length HA10-ADAMTS13D complex in solution and three-dimensional modeling.** **a**, The weight concentration distribution $c(s_{20,w})$ as a function of sedimentation coefficient $s_{20,w}$ obtained by analytical ultracentrifugation (AUC) for ADAMTS13D, full-length HA10, and mix solutions of full-length HA10 + ADAMTS13D with various mixing molar ratios. **b,** Mixing molar ratio dependence of the $s_{20,w}$-value corresponding to full-length HA10 and full-length HA10-ADAMTS13D complex. Peak $s_{20,w}$-values were determined by Gaussian fitting of $c(s_{20,w})$, and the error bars represent the standard errors of the fitted peak positions. The solid lines represent the mixing-ratio dependence of the weight-average sedimentation coefficient for full-length HA10 and the full-length HA10–ADAMTS13D complex, calculated using the method of the Dam and Schuck method[36]. The black, magenta, yellow, and cyan lines correspond to calculated weight-average sedimentation coefficients for dissociation constant $K_d$ values of 1 nM, 10 nM, 100 nM, and 1 μM, respectively. Because the calculated curves become relatively insensitive to $K_d$ value in the submicromolar range, the $K_d$ value could not be determined precisely from the AUC data alone. Nevertheless, the experimental weight-average sedimentation coefficients were better reproduced by the curves calculated for submicromolar $K_d$ values, supporting a relatively high-affinity interaction. At mixing molar ratios above 1:2, the sedimentation coefficient saturated at 7.27 S, corresponding to a complex with a binding stoichiometry of 1:2. **c**, Red circles show the experimental SAXS profile for a full-length HA10-ADAMTS13D complex with a binding stoichiometry of 1:2. Black line represents the calculated scattering profile for the full-length HA10-ADAMTS13D complex model (Fig. 2d). The scattering intensity is normalized by the concentration. The residuals between the experimental and calculated scattering profiles are shown in the lower panels. **d**, Partial structural analysis of the full-length HA10 in full-length HA10-ADAMTS13D complex using inversed contrast matching SANS. Black and red circles represent the scattering profile of hydrogenated (h)-full-length HA10 in 100%$D_2O$ and mix solution of (h)-full-length HA10 and 75% deuterated (75d)-ADAMTS13D in 100%$D_2O$, respectively. The scattering intensities are normalized by concentration of (h)-full-length HA10 ($c_{HA10}$). Because the scattering length density of (75d)-ADAMTS13D matches that of 100% $D_2O$, (75d)-ADAMTS13D is invisible to neutron scattering. The agreement between the two scattering profiles means that the partial structure of full-length HA10 in the complex is identical to full-length HA10 alone in solution. Blue solid line represents the calculated scattering profile for full-length HA10 in HA10-ADAMTS13D complex. The residuals between the experimental and calculated scattering profiles are shown in the lower panels.

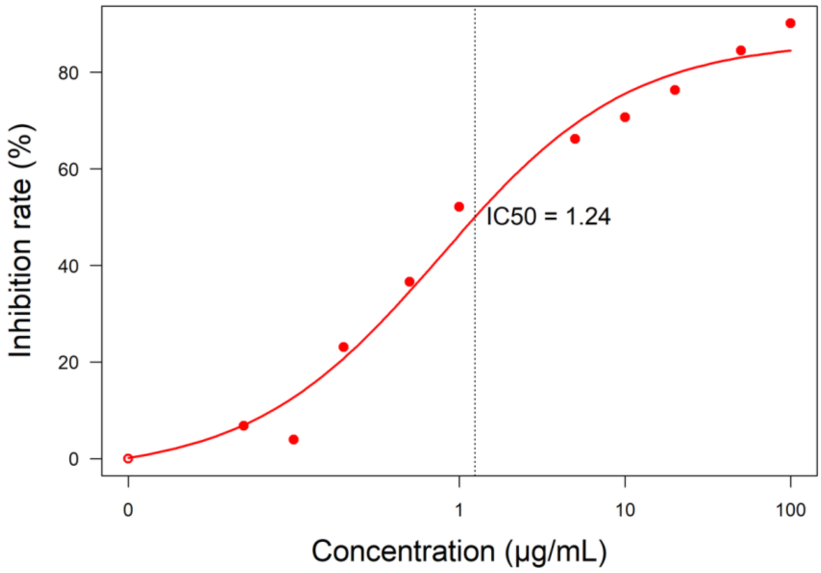


| HA10 concentration (μg/mL) | ADAMTS13 activity(%) |
|---|---|
| 0 | 71 |
| 1 | 34 |
| 10 | 20.8 |
| 100 | 7 |

**Extended Data Fig. 5: Concentration–response curve of HA10 in vitro.** The concentration-dependent ADAMTS13 inhibitory effect of HA10 on ADAMTS13 activity was determined by incubating standard human plasma with 100 μg/mL to 0.05 μg/mL. The values at dose 0 control was also measured to calculate the inhibitory rate. The $IC_{50}$ value was estimated by using R version 4.3.2 packages.

Abbreviations: ADAMTS13, a disintegrin and metalloproteinase with thrombospondin type 1 motif 13; IC50, half maximal inhibitory concentration.

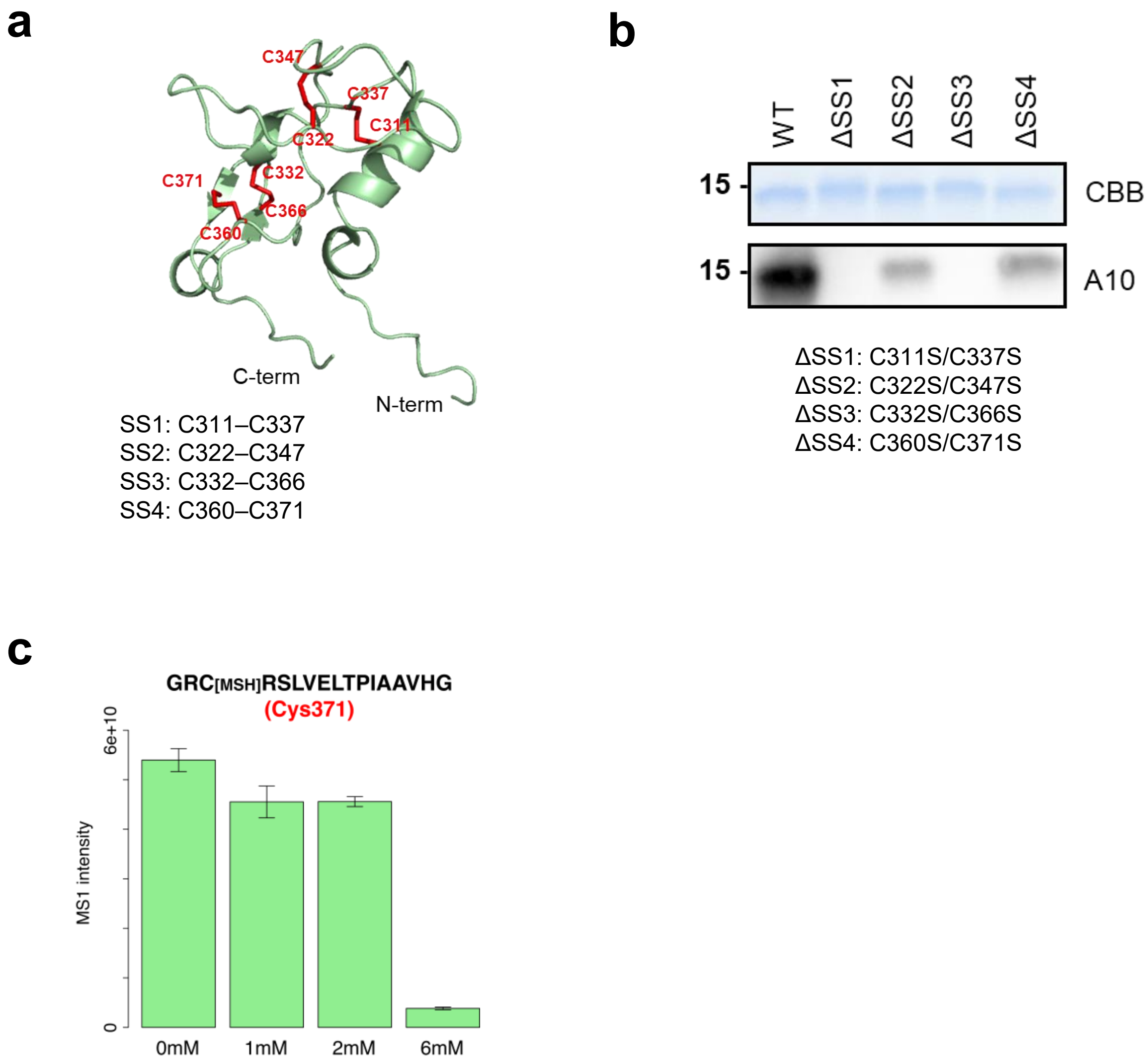


**Extended Data Fig. 6: Disulfide bond organization and GSH-responsive cysteine accessibility of ADAMTS13D. a**, Structural model of ADAMTS13D showing the four intramolecular disulfide bonds highlighted in red. **b**, mA10-binding assays of ADAMTS13D ΔSS mutants in which individual Cys-Cys pairs were substituted with Ser-Ser pair. **c**, Mass spectrometric analysis of GSH-induced thiol–disulfide exchange in ADAMTS13D. The abundance of the C371-containing peptide was quantified after incubation with increasing concentrations of GSH.

```
Human       290 PGSAGHPPDAQPGLYYSANEQCRVAFGPKAVACTFAREHLDMCQALSCHTDPL 342
Cynomolgus  294 PGSAGRQPEAQPGLYYSANEQCRVAFGPKAVACTFSREHLDMCQALSCHTDPL 346
Porcine     295 SWAAGRPPEAHPGLYYGADEQCRVAFGPTAVACTFAREQLDMCQALSCHTDPL 347
Bovine      295 SGPAGQPPEVQPGLYYGADEQCRVAFGPTAVACTFRGEHLDMCQALSCHTDPL 347
Caprine     295 PGPAGRPPEAQPGLYYGADEQCRVAFGPTAVACTFRGEHLDMCQALSCHIDPL 347

                      ▼▼ ▼
Human       343 DQSSCSRLLVPLLDGTECGVEKWCSKGRCRSLVELTPIAAVHG 385
Cynomolgus  347 DQSSCSRLLVPLLDGTECGVEKWCSKGRCRSLVELTPIAAVHG 389
Porcine     348 DQSSCSRLLIPLLDGTECGVGKWCSKGHCRSLAELAPVGAVHG 390
Bovine      348 DPSSCSRLLIPLLDGTECGVGKWCSKGHCRSLAELAPVGVVHG 390
Caprine     348 DPSSCSRLLIPLLDGTECGVEKWCSRGHCRSLAELAPVGVVHG 390
```

**Extended Data Fig. 7: Multiple sequence alignment of the disintegrin-like domain of ADAMTS13 from different species.** Red arrows indicate the conserved exosite residues (R349, L350, and V352) known to anchor the unfolded VWF A2 domain.

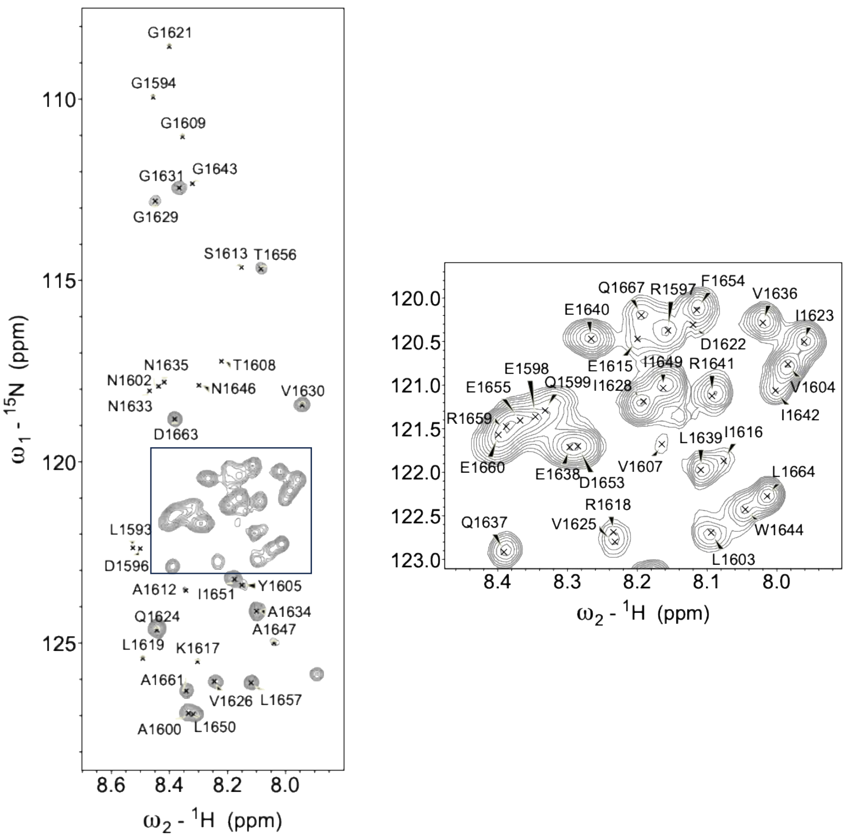


**Extended Data Fig. 8: NMR signal assignment of stable isotope-labeled VWF73.**

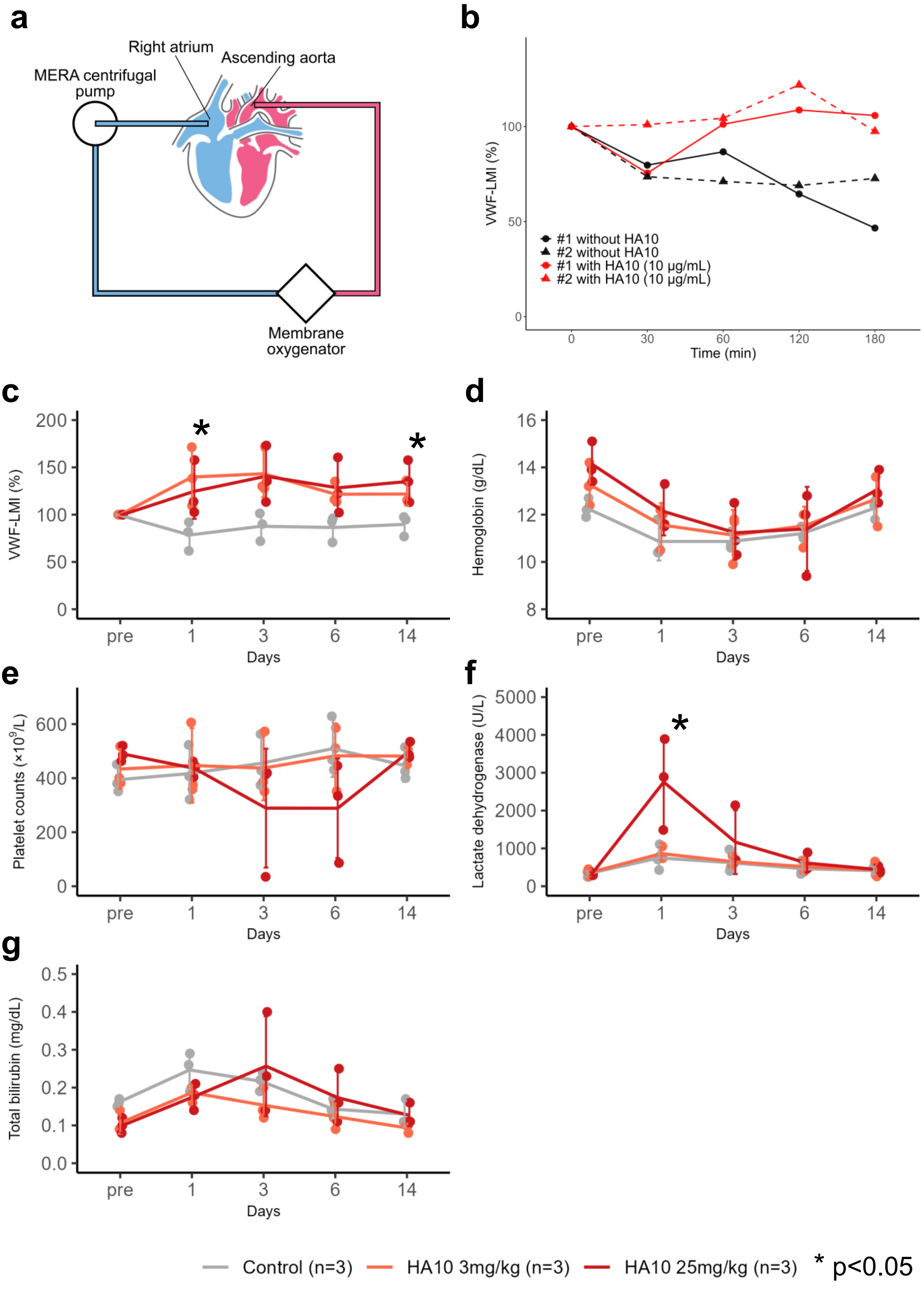
a
Right atrium
Ascending aorta
MERA centrifugal pump
Membrane oxygenator
b
VWF-LMI (%)
Time (min)
#1 without HA10
#2 without HA10
#1 with HA10 (10 μg/mL)
#2 with HA10 (10 μg/mL)
c
VWF-LMI (%)
Days
d
Hemoglobin (g/dL)
Days
e
Platelet counts (×10^9/L)
Days
f
Lactate dehydrogenase (U/L)
Days
g
Total bilirubin (mg/dL)
Days
Control (n=3)
HA10 3mg/kg (n=3)
HA10 25mg/kg (n=3)
* p<0.05

**Extended Data Fig. 9: Cynomolgus monkey studies of the AVWS model and GLP-grade HA10 safety evaluation.** **a–b**, Cynomolgus monkey AVWS model. **a**, Schematic illustration of the cynomolgus monkey AVWS model. **b**, Temporal changes in VWF-LMI in the PCPS-implanted cynomolgus monkey AVWS model. Error bars represent mean ± SD. **c–g**, Temporal changes in laboratory parameters following single-dose administration of GLP-grade HA10, comparing the 25 mg/kg group (n=3), the 3 mg/kg group (n=3), and the control group (n=3). **c**, VWF-LMI. **d**, hemoglobin. **e**, platelet counts. **f**, lactate dehydrogenase. **g**, total bilirubin. Error bars represent mean ± SD. *$p < 0.05$. Abbreviations: VWF, von Willebrand factor; VWF-LMI, VWF large multimer index.
